# CK2 directly controls CARD9 protein homeostasis

**DOI:** 10.1101/2025.02.03.636288

**Authors:** Aya M. Kelly, Leidy C. Merselis, Benjamin Causton, Olivia K. Calder, Hoson Chao, Margaret Prack, Laurel B. Stine, Yuka Amako, Tao Wang, Ali E. Yesilkanal, Kendall J. Condon, Meekyum O. Pierce, Scott H. Watterson, Jason Guernon, Amit Anand, Sabariya Selvam, Sayali Dagde, Emily Holzinger, Lauren Piasecki, Travis A. Pemberton, Emily Fu, McKenna Joint, Hayden Sandt, Ashish K. Singh, Tai Wang, Denise Cobb, Xiaochun Ni, Geraint H. M. Davies, Alan F. Corin, Sophie Roy, Stephen C. Wilson

## Abstract

CARD9 is an attractive target for therapeutic intervention because the human genetic data provides strong evidence for the causal role of CARD9 in both protection and susceptibility to autoimmune disease. Expression quantitative trait loci (eQTLs) link higher CARD9 expression to increased disease risk and lower CARD9 expression to protection. Additionally, a rare allele variant leading to a C-terminal truncation of CARD9 (CARD9Δ11) and subsequent loss-of-function is also protective for inflammatory bowel disease (IBD). The mechanism of CARD9-driven inflammation through scaffold assembly with BCL10 and MALT1 (CBM complex) suggests a durable inflammatory signal driven by increasing levels of CARD9. Therefore, CARD9 depletion is a desired therapeutic strategy for drug discovery, yet a difficult one due to the nature of CARD9 as an adaptor protein target and the limited number of chemical tools available to engage it. Here, we uncover through a protein homeostasis screen that casein kinase 2 (CK2/CSNK2) inhibition leads to cellular CARD9 depletion. Following up with arrayed CRISPR screening, we identify key casein kinase isoforms/subunits responsible for CARD9 depletion. We find that CK2 directly binds to CARD9 and phosphorylates S424/S425 as well as S483/S484. Orthosteric CK2 inhibition prevents CK2 binding to CARD9 and leads to CARD9 destabilization. We show that the interaction between CK2 and CARD9Δ11 is significantly attenuated and not sensitive to CK2-mediated protein stabilization. The CK2-driven CARD9 depletion mechanism is preserved outside of immortalized cell lines and conserved between primary, differentiated mouse and human dendritic cells. Finally, we demonstrate therapeutic proof of concept *in vivo* using CK2 inhibition to deplete CARD9 in murine whole blood, peritoneum, and colon. Our study expands the scope of cellular consequences dealt by kinase inhibition, offers an unconventional approach for engaging a therapeutically intractable target, and identifies a novel mechanism that could contribute to disease protection conferred by the CARD9Δ11 allele.

## INTRODUCTION

Caspase recruitment domain-containing protein 9 (CARD9) is an adaptor protein that plays a critical role in the innate immune response, particularly in the signaling pathways initiated by C-type lectin receptors (CLRs) such as Dectin-1.^1^ CARD9 is predominantly expressed in myeloid cells, and its activation forms higher-order protein complexes with BCL10 and MALT1 upon phosphorylation by PKCδ, leading to NF-κB induction.^2^ The cytokine response driven by the CARD9 pathway is essential for mounting effective immune responses against fungal, bacterial, and viral pathogens. However, in recent years CARD9 has garnered significant attention due to its association with several inflammatory diseases based on human genetics.

Human genetic studies provide unique insights into the causal role of CARD9 in immune disease pathology based on three distinct genetic signals (Fig. 1a). First, genome-wide association studies (GWAS) identified a low frequency splicing variant (rs141992399) that results in the exclusion of exon 11 (∼0.3% population allele frequency).^3^ This variant has a strong protective effect with ∼4-fold decrease in risk of IBD for each allele (odds ratio = 0.29, *p*= 1e^−16^). One study found evidence that the resulting variant transcript, CARD9Δ11, causes a C-terminal truncation and may prevent downstream activation, implicating CARD9 loss-of-function (LoF) in disease protection.^4^ Second, a common variant near CARD9 is associated with increased IBD risk and increased CARD9 expression in whole blood and immune cells, including monocytes (naïve and stimulated), neutrophils, and CD4^+^ T-cells.^5^ Interestingly, CARD9Δ11 eliminates the risk conferred by this signal when the splice variant and common variant are on the same haplotype, underscoring the magnitude of protection conferred by CARD9 LoF.^3^ GWAS fine-mapping identified an additional intronic signal in CARD9 that was specific to Crohn’s Disease (CD). In mechanistic agreement with the other two signals, the lead variant is associated with CD protection and decreased CARD9 expression in monocytes.^6^

**Fig. 1.**
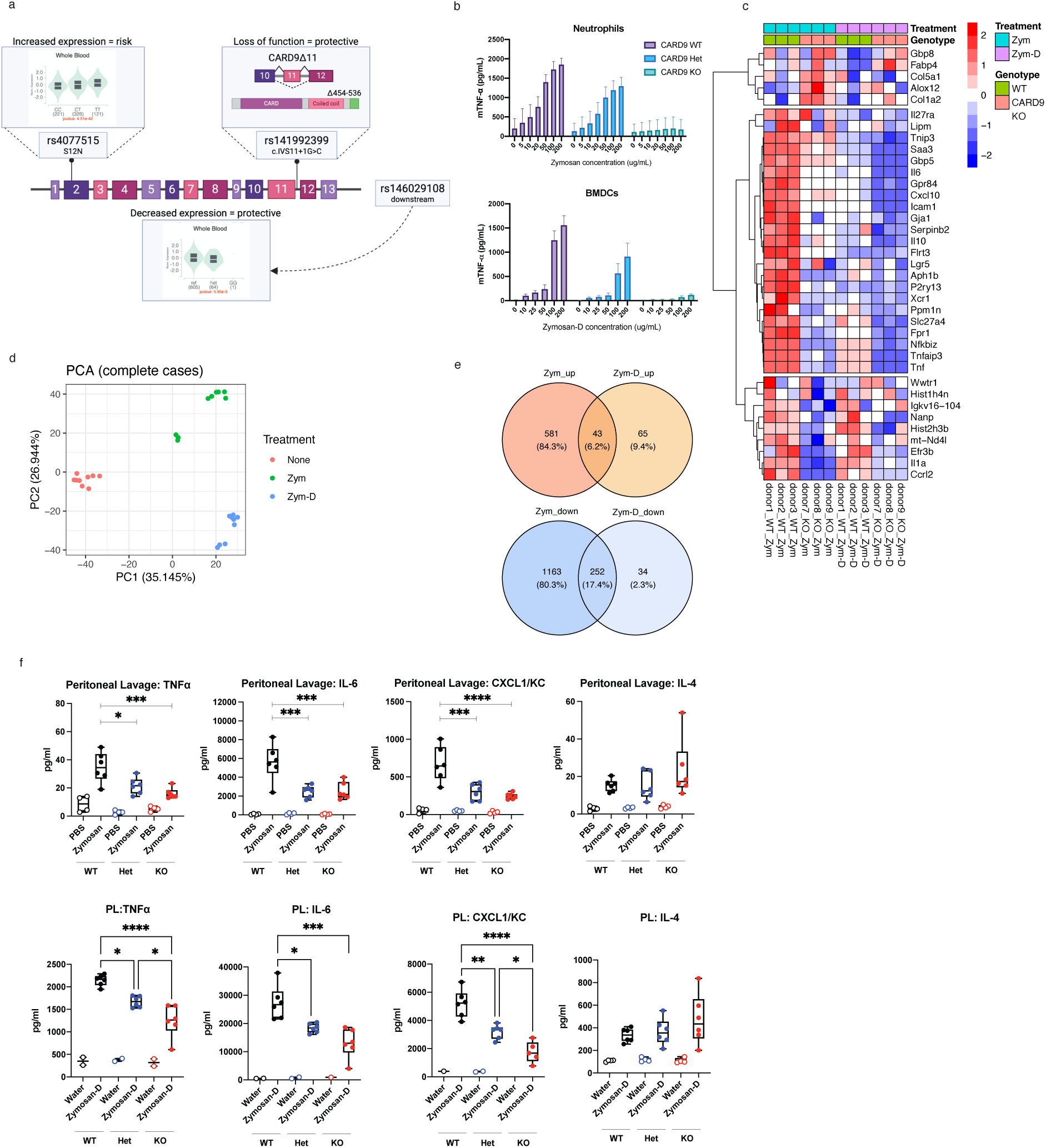
CARD9 depletion supports anti-inflammatory directionality provided by human genetics. **(a)** Three genetic variants with independent effects on CARD9 function and IBD risk. Locations in reference to the CARD9 gene structure. Clockwise: rs4077515-T is a common missense variant (S12N) associated with increased IBD risk and increased CARD9 expression in whole blood and monocytes (violin plot for whole blood in GTEx shown); rs141992399 is a lower frequency splice variant CARD9 loss-of-function and decreased risk of IBD; rs146029108 is a common intergenic variant associated with decreased risk of IBD and decreased CARD9 expression in whole blood and monocytes (violin plot for whole blood in GTEx shown). **(b)** TNF-α response in neutrophils (5-hour stimulation) and BMDCs (overnight stimulation) by zymosan or zymosan-D stimulation is dependent on CARD9 levels (n = 3). Data represented as mean with SD in graph. **(c)** Heatmap shows expression changes in top differentially expressed genes upon zym and zym-D treatment (2-hour stimulation) in WT and CARD9 KO samples at q-value threshold of 0.01 and log2-fold change cutoff of 2. Expressed values for each gene upon zym/zym-D treatment is normalized to the baseline expression level in the no-treatment sample from the same donor, and z-transformed. **(d)** Principal component analysis of experimental samples based on gene expression data after removing unknown confounding effects by surrogate variable analysis. **(e)** Venn diagrams show the overlap between the targets upregulated (top) or downregulated (bottom) by zym and zym-D. Percentages of unique and common targets are indicated in parentheses. **(f)** Luminex levels of TNFα, IL-6, CXCL1/KC and IL-4 in the peritoneal lavage (PL) fluid of either PBS (n = 4), zymosan (50 mg/kg, n = 6), water (n = 4), or zymosan-D (50 mg/kg, n = 6)-treated WT, CARD9^+/−^ heterozygous or CARD9^−/−^ KO mice. Statistical significance determined by one-way ANOVA with adjusted p value depicted (*-*p* ≤ 0.05, **-*p* ≤ 0.01, ***-*p* ≤ 0.001, ****-*p* ≤ 0.0001).

Based on the human genetics, we hypothesized that pharmacological depletion of CARD9 could be protective for autoimmune disease. However, little is known about mechanisms that regulate CARD9 expression or stability. TRIM62 has been reported to ubiquitinate CARD9 between its CARD domain and first coiled-coiled domain, leading to its activation.^4^ Conversely, USP15^7^ and OTUD1^8^ have been described to deubiquitinate CARD9 and reported to attenuate and potentiate CARD9-mediated signaling, respectively. Nonetheless, all studies focus on subsequent CARD9 activation/deactivation upon ubiquitination or de-ubiquitination events, not changes in CARD9 expression. As such, we performed a protein homeostasis screen to identify pharmacological modulators of CARD9 expression with a focus on CARD9 depletion.

From this screen we discover from a small molecule protein homeostasis screen that CK2 (casein kinase 2, CSNK2) inhibition leads to cellular CARD9 depletion and that it is a conserved mechanism among cancer cells, human primary cells, and mouse primary cells. We find that CK2 binds directly to the C-terminus of CARD9 and phosphorylates CARD9 at S424/S425 and S483/S484. CK2 inhibition effectively disrupts the protein-protein interaction between CARD9 and CK2, leading to CARD9 depletion in the cell. Finally, we leverage this phenomenon to demonstrate therapeutic proof of concept to deplete CARD9 in various compartments within a zymosan-treated mouse.

## RESULTS

### Experimental Modeling of CARD9 Depletion Supports Anti-Inflammatory Directionality Provided by Human Genetics

CARD9 deficiency has been associated with both increased and decreased susceptibility to autoimmunity. On the one hand, CARD9 deficiency in mice is linked to impaired surveillance and response to maladaptive gut flora, leading to their burgeoning and subsequent induction of colitis.^9–11^ On the other hand, CARD9 loss has been shown to suppress *Malassezia restricta-* exacerbated dextran sulfate sodium (DSS) induced colitis in mice as well as attenuate autoantibody-induced arthritis and dermatitis.^12,13^ Since human genetics suggests CARD9 deficiency and LoF are protective against disease (Fig. 1a), we focused on evaluating the effects of CARD9 loss *in vitro* and *in vivo*.

It is well-described that myeloid-derived cells from CARD9 KO mice have attenuated induction of proinflammatory cytokines upon zymosan stimulation *in vitro*.^1^ We confirmed bone marrow-derived dendritic cells (BMDCs) and neutrophils from CARD9^+/−^ and CARD9^−/−^ mice exhibit reduced TNFα response upon zymosan stimulation (Fig. 1b, Supp. Fig 1). However, zymosan signals through both Dectin-1 and TLR2, leading to mixed pathway contributions that drive pro-inflammatory signaling. Hot alkali treated zymosan (zymosan-D) has been reported to dissociate the TLR2 signaling effects from zymosan, while preserving Dectin-1 signaling.^14^ In our hands, zymosan-D stimulation demonstrated greater dependence on CARD9 in BMDCs than zymosan stimulation (Fig. 1b, Supp. Fig 1). Nonetheless, a transcriptional response comparison between zymosan and zymosan-D has not been reported to our knowledge. Given that CARD9 signaling specifically in neutrophils has been directly linked to autoimmune pathophysiology in K/BxN serum treated mice^12^ as well as SKG mice^15^, we investigated the effects of CARD9 signaling on neutrophils upon zymosan and zymosan-D stimulation using RNA-seq.

Both zymosan and zymosan-D elicited CARD9-dependent proinflammatory programming as evident by *Tnf*, *Tnfaip3*, *Il-6*, *Il-1*α, and *Cxcl10* in the transcriptional heatmap (Fig. 1c). Pathway analysis illustrates this commonality with shared annotations for “inflammatory response” and “response to molecule of bacterial origin” that are suppressed in both conditions (Supp. Fig. 2). However, neutrophils treated with zymosan and zymosan-D clustered distinctly from each other based on principal component analysis (PCA), indicating differentiation from each other (Fig. 1d). Between zymosan and zymosan-D only 6.2% and 17.4% of genes upregulated and downregulated were shared, respectively. In CARD9 KOs, zymosan uniquely elicited pathways annotated for “DNA repair”, “DNA replication”, “spindle assembly”, “chromosome organization” and others that indicate a strong association with cellular division and DNA maintenance (Supp. Fig. 2). We also found that the breadth of elicited cytokines that are CARD9-dependent are significantly larger for zymosan versus zymosan-D (Supp. Fig. 3). Since zymosan and zymosan-D demonstrated both shared and disparate traits between each other, we decided to evaluate their CARD9 dependence in vivo within a peritonitis model.

It has previously been reported that zymosan-induced peritonitis is exacerbated in CARD9^−/−^ mice.^16^ While this result is inconsistent with the mechanism of action associated with CARD9, it is testimony to the bidirectionality of CARD9 described above and found throughout the literature. From our own studies, we found that CARD9^+/−^ and CARD9^−/−^ mice elicited cytokine protection within either a zymosan or zymosan-D peritonitis model (Fig. 1f, Supp. Fig. 4). Notably, TNFα, IL-6, KC, and other CARD9-dependent mediators demonstrated consistent and significant attenuation in both the peritoneal lavage and serum (Fig. 1f, Supp. Fig. 4). These data support the therapeutic hypothesis that CARD9 depletion could attenuate pro-inflammatory cytokine signaling associated with autoimmune disease.

### Protein homeostasis screen for CARD9 depletion reveals connection with CK2

To support a high-throughput screen (HTS) for the identification of small molecule cellular depleters of CARD9, an endogenously edited THP-1 cell line containing an N-terminus HiBiT tag on CARD9 was engineered using CRISPR/cas9 (Fig 2a). During subsequent screen triage, we found that ∼95% of our hits came from chemical matter associated with internal CK1 or CK2 programs (CK#), suggesting a biological mechanism of action between CARD9 and CK# (Fig. 2b). Inhibitors from the CK1 program (BMS-321), CK2 program (BMS-926, BMS-713, and BMS-747)^17–19^, and a clinical CK2 inhibitor (CX4945, silmitasertib) were chosen as representatives for study based on potency and PK profile (Fig. 2c). High affinity CK2 inhibitors (K_D_ ≤ 1 nM) demonstrated excellent cellular potencies for CARD9 depletion (DC_50_ <25 nM), leading to >50% CARD9 depletion in wild type (WT) THP-1s by AlphaLISA (Fig. 2d). CARD9 depletion was validated by global proteomics (Supp. Fig 5), and we found that our internal series demonstrated a significant correlation (R = 0.65) between CARD9 depletion and cellular CK2 inhibition as measured by growth rate attenuation in the CK2-dependent SNCU-1 cell line (Fig. 2e). While CARD9 depletion from CX4945 was markedly less potent (i.e. ∼4 μM), this is consistent with its reported cellular potency^20^ compared to the low nanomolar cellular potency of our internal series.^17^ Since CX4945 has been reported to inhibit a few off-target kinases (e.g. GSK3β and DYRK1A),^21^ we sought to confirm that the CARD9 depletion-induced by CK2 inhibition was not due to off-target pharmacology.

**Fig. 2.**
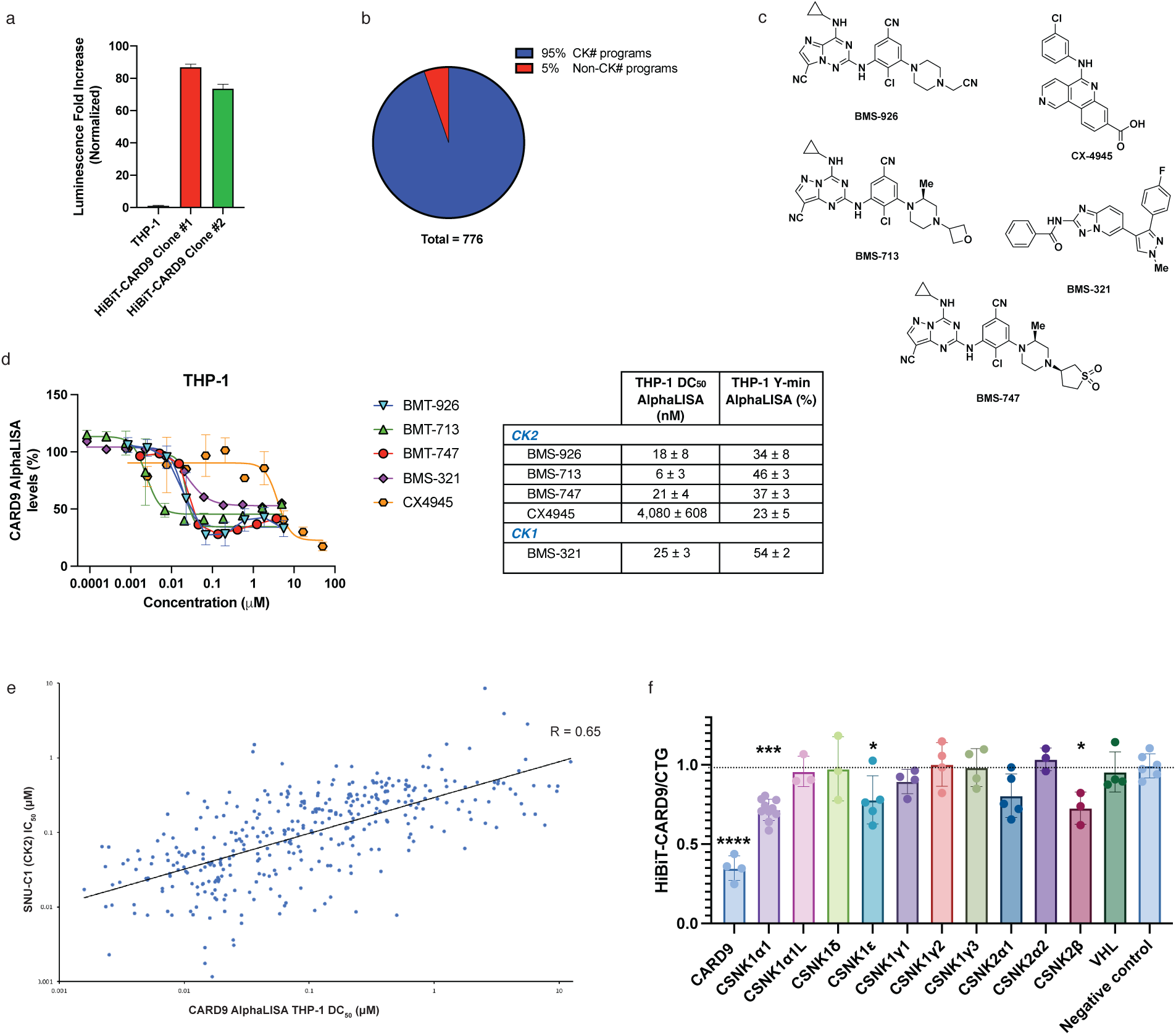
Discovery of CARD9-CK2 interaction. **(a)** Luminescence of THP-1 HiBiT-CARD9 constructs used for screening. Data represented as mean with SD in graph. **(b)** THP-1 CARD9 HiBiT HTS identified CARD9 depleters highly enriched with known CK# compounds. **(c)** Chemical structures of CK1 compound (BMS-321) and CK2 compounds (BMS-926, BMS-713, BMS-747, and CX4945). **(d)** CK# inhibition leads to CARD9 depletion in THP-1s as measured by CARD9 AlphaLISA (overnight incubation, n = 3). Data represented as mean with SD in graph. **(e)** Correlation between CK2 inhibition in a SNU-C1 functional assay and CARD9 depletion. **(f)** Impacts of CK# loss with arrayed CRISPR sgRNAs on HiBiT-CARD9 levels normalized to cell viability as measured by CellTiter-Glo® (CTG). Significant decreases in normalized HiBiT-CARD9 were observed for sgRNAs against CARD9 as a control, CSNK1α1, CSNK1ε, and CSNK2β. Data represented as mean with SD in graph. Statistical significance determined by one-way ANOVA with adjusted p value depicted (*-*p* ≤ 0.05, **-*p* ≤ 0.01, ***-*p* ≤ 0.001, ****-*p* ≤ 0.0001).

By leveraging our THP-1 HiBiT-CARD9 cell line, we executed an arrayed CRISPR/cas9 screen to confirm that CARD9 depletion was not due to off target pharmacology associated with CK# inhibitors as well as evaluate the effect of knocking out specific CK1 and CK2 subunits on cellular CARD9 levels (Fig. 2f). CK1 is characterized as family of multiple kinase isoforms (i.e. CSNK1α, CSNK1β, CSNK1ε, etc.) that act separately and discretely from each other, whereas CK2 forms a larger complex comprised of one CSNK2α1 subunit, one CSNK2α2 subunit, and two CSNK2β subunits.^22^ CSNK2α1 and CSNK2α2 are enzymatic components of the CK2 complex that are directly targeted by small molecule inhibition, whereas the CSNK2β subunit is regulatory in nature. CARD9 protein levels showed significant dependence on CSNK1α1, CSNK1ε, and CSNK2β, supporting that CARD9 depletion from CK# inhibition was not due to pharmacological off target effects. It is worth noting that a near significant dependence on CSNK2α1 (*p* = 0.067) was also observed. These data validated a direct or indirect interaction between CK# and CARD9.

To date there is only one reference in the literature on both CARD9 and CK2 and none on both CARD9 and CK1. It has been reported that pVHL (Von Hippel-Lindau tumor suppressor) acts as an adaptor protein between CARD9 and CK2 in cancer cells, which promotes indirect phosphorylation of CARD9 by CK2.^23^ pVHL is a well-described component of E3 ubiquitin ligase machinery that targets HIF1α for protein degradation.^24^ Interestingly, pVHL was reported not to target CARD9 for destruction in this context.^23^ In accord, we found that knocking out pVHL had no significant effect on cellular CARD9 protein levels (Fig. 2f), suggesting an alternative and unique mechanism for CK2-dependent regulation on CARD9 protein levels that does not require VHL as the adaptor protein.

### CK2 inhibition depletes CARD9 in human primary dendritic cells

Since our data collected thus far only indicated a role for CK2 inhibition in CARD9 depletion within cancer cell lines, we investigated whether this effect was generalizable to human primary cells. We found that CK2 inhibition with our chemical series exhibited comparable cellular potencies for CARD9 depletion (DC_50_ <50 nM) and levels of CARD9 depletion (Y_min_ <50%) in human primary dendritic cells across multiple donors as found in THP-1 cells (Fig. 3a, 3b). CX4945 also exhibited similar micromolar potency in human primary dendritic cells (∼2 μM) compared to THP-1 cells (∼4 μM). No significant toxicity was observed for any of the compounds as measured by lactate dehydrogenase (LDH) assay (Fig. 3c). However, we found that the CK1 inhibitor, BMS-321, did not deplete CARD9 in human primary cells, suggesting that the CARD9 depletion mechanism by CK1 is unique to immortalized cell lines. Since the CARD9-CK1 mechanism was not conserved in human primary cells, we decided to focus our attention on elucidating the mechanism for CK2-driven CARD9 depletion.

**Fig. 3.**
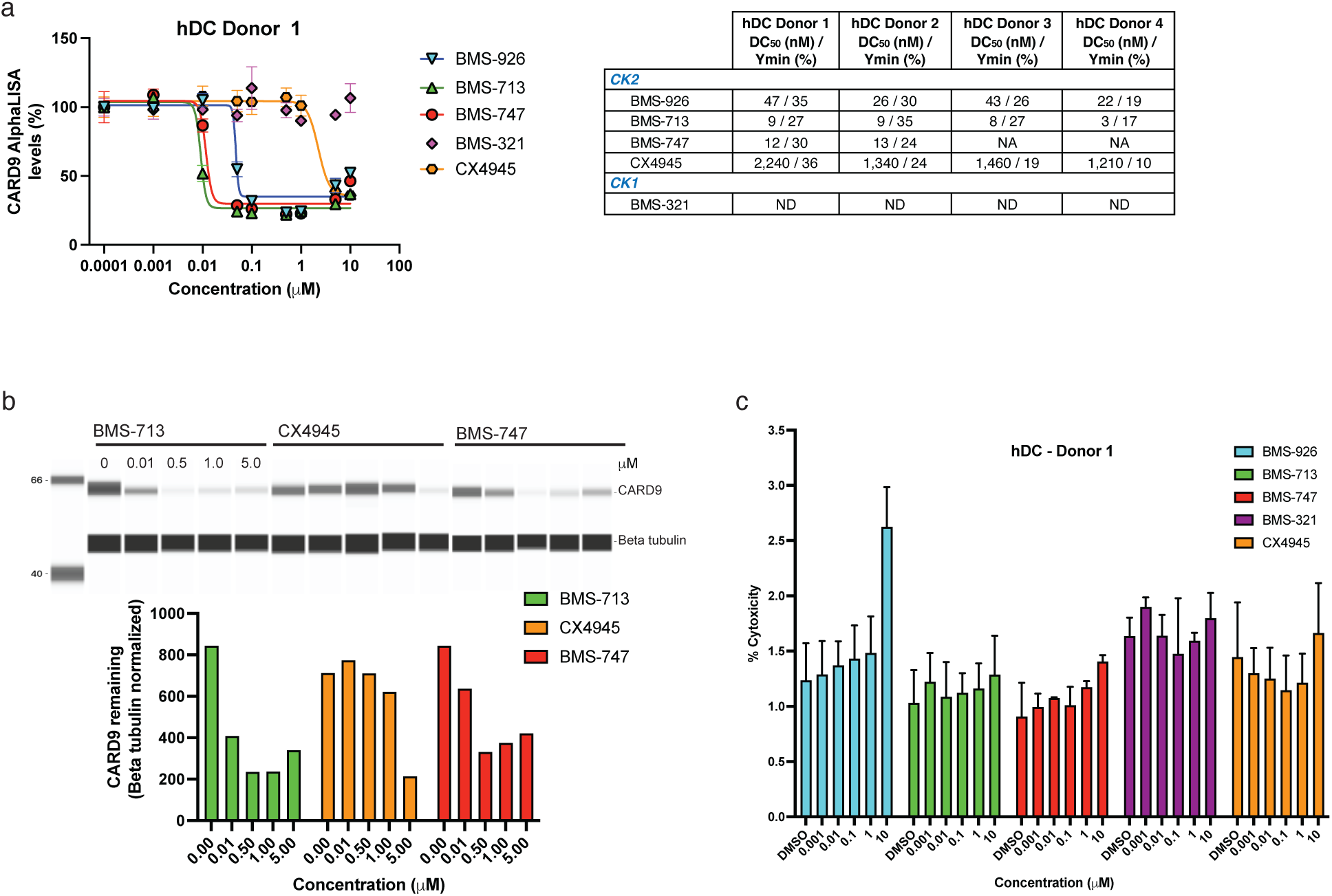
CK2 inhibition elicits CARD9 depletion in differentiated human dendritic cells. **(a)** CK2 inhibition leads to CARD9 depletion in human primary dendritic cells (hDCs) as measured by CARD9 AlphaLISA (overnight incubation). Data represented as mean with SD in graph. **(b)** Jess western blot of CARD9 depletion in hDCs. **(c)** LDH cytotoxicity analysis in hDCs (n = 3). Data represented as mean with SD in graph.

### CK2 binds directly to CARD9 and phosphorylates CARD9 at S424/S425 and S483/S484

It is well-documented that CK2 can influence transcription by phosphorylating transcription factors and components of the transcriptional machinery.^25^ To rule out transcription as the CK2-dependent driver of CARD9 depletion, we performed RT-qPCR at a 6 hour time point in THP-1s. We found that our CK2 chemical inhibitors did not result in CARD9 mRNA depletion and in fact, elicited a modest elevation of CARD9 transcript (Fig. 4a) that did not track with the concentration response curve (CRC) as shown in Fig. 2d. Interestingly, we did observe a significant level of CARD9 transcript depletion that correlated with the potency of our CK1 inhibitor, BMS-321, suggesting that the CARD9-CK1 mechanism in immortalized cells has a transcriptional component (Fig. 4a).

**Fig. 4.**
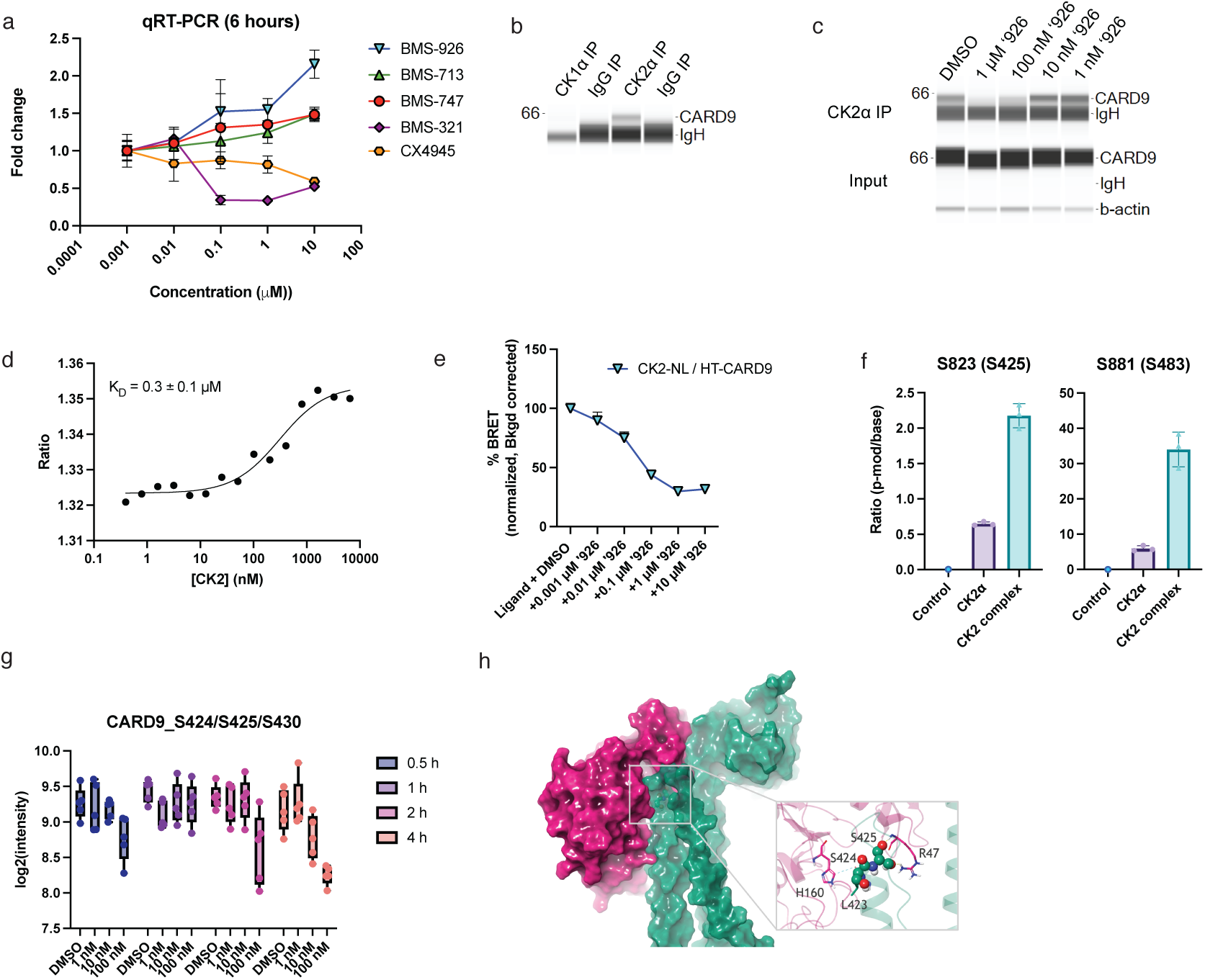
CK2 binds directly to CARD9 and phosphorylates S424/S425 and S483/S484. **(a)** qRT-PCR in THP-1s of CK# inhibitors with CARD9 probe (n = 3). Data represented as mean with SEM in graph. **(b)** co-IP of 3x FLAG-CK1α and CK2α blotted for CARD9 and visualized by JESS Western blot. **(c)** CK2-CARD9 interaction is disrupted by BMS-926 in a dose-dependent fashion (1-hour incubation). **(d)** Spectral shift (650/670 nm) dose-response curve demonstrating binding of a fluorescently labelled MBP-CARD9 to CK2. K_D_ reported as mean ± SD (n = 3). **(e)** NanoBRET dose-dependent disruption of CK2-CARD9 protein-protein interaction by a CK2 inhibitor at 4 hours. Data points reported as mean ± SD (n = 4). **(f)** Abundance ratio of the monophosphorylated peptide over unmodified peptide from MBP-CARD9 containing the indicated serine residue. **(g)** Intensity of the monophosphorylated peptide CARD9 aa416-432 at different time points and dose of BMS-713. **(h)** Surface representation of protein-protein docked pose of CARD9 (green) and CK2 (pink). The observed phosphorylation site of CARD9 inserted in the ATP binding pocket of CK2 and the complex is stabilized via pi-cation interaction between L423 (CARD9) and H160 (CK2), and H-bonds between S425 (CARD9) and R47 (CK2).

We next examined whether CK2-driven CARD9 depletion was mediated by a protein-protein interaction. We found that CARD9 co-immunoprecipitated with CSNK2α1 (Fig. 4b) and could be competed off with titration of BMS-926 (Fig. 4c). Binding assays using spectral shift with the full CK2 complex and in vitro purified MBP-CARD9 afforded a K_D_ of 0.3 ± 0.1 μM, indicating a direct protein-protein interaction (Fig. 4d). To further validate a direct cellular interaction, we utilized a cellular nanoBRET assay with CSNK2α1-NanoLuc and HaloTag-CARD9, which led to a significant proximity-driven BRET signal that attenuated with titration of BMS-926 (Fig. 4e).

CK2 is a highly pleiotropic kinase and phosphorylates hundreds of protein substrates.^26^ It has previously been reported that CK2 and CARD9 are both recruited to pVHL, where pVHL mediates CARD9 hyper-phosphorylation by CK2. However, CARD9 was transiently transfected in this study, leading to non-physiologically relevant phosphorylation driven by the higher concentrations of CARD9 achieved after transient transfection.^23^ In our hands, incubation of MBP-CARD9 protein with the CK2 complex or CSNK2α1 enzyme in the absence of VHL led to significant phosphorylation at distinct sites on CARD9 including: S424, S425, S483, and S484 (Fig. 4f, Supp. Fig. 6). Mass spectrometry captured 31 out of 33 serines and 16 out of 17 threonines from CARD9. To strengthen these results, we incubated THP-1s with our CK2 inhibitor and monitored CARD9 phosphorylation over time by mass spectrometry. Cellular phosphoproteomics identified 9 serines and 1 threonine associated with CARD9. One mono-phosphorylated peptide containing S424/S425/T431 showed significant downregulation upon CK2 inhibitor treatment by 4 hours (Fig. 4g) and was also in accord with the relative potency of our CK2 inhibitors (Fig. 2d).

Given the observed protein interaction between CSNK2α1 and CARD9, we sought to structurally model a snapshot of the phosphorylation event. Due to the lack of publicly available full-length structure of CARD9 to date, the predicted monomeric structure of CARD9 from AlphaFold^27^ was used along with the kinase domain of CK2 (PDB: 5ZN1) in protein-protein docking, where distance constraints between CK2 and S424/S425 of CARD9 were applied given the observed phosphorylation pattern. Among the top three protein-protein docked conformations, we found that one pose is significantly more stable than the other two based on the root mean square deviation and root mean square fluctuation from molecular dynamics simulations of modeled conformations.

In the top, most stable predicted pose of CK2 and CARD9, we see that the local region surrounding S424/S425 of CARD9 is inserted in the ATP binding pocket of CK2 (Fig. 4h). The CARD9:CK2 complex is stabilized via stable pi-cation interaction between H160 of CK2, located just outside of the ATP binding site, and backbone carbonyl of L423, the residue immediately preceding S424, of CARD9. In addition, there are several stable H-bonds between K44-Q127, R47-S425, R47-Q131, K49-D426 (CK2-CARD9 pairs), all of which are near the P-loop covering the ATP binding pocket of CK2, resulting in the stable protein-protein interface that is likely contributing to the phosphorylation event of CARD9 by CK2.

### CARD9 phosphomimetic screen identifies activating CARD9 T231D mutation

Since CARD9 is phosphorylated by CK2, we aimed to understand how phosphorylation events on CARD9 (Fig. 5a) could affect CARD9-dependent NF-κB signaling. As such, we performed a phosphomimetic screen of CARD9 in HEK293T using transient transfection of CARD9, BCL10 and an NF-κB dual-luciferase-based system. The screen covered 84% of all serines (28 out 33) and 71% of all threonines (12 out of 17) on CARD9.

**Fig. 5.**
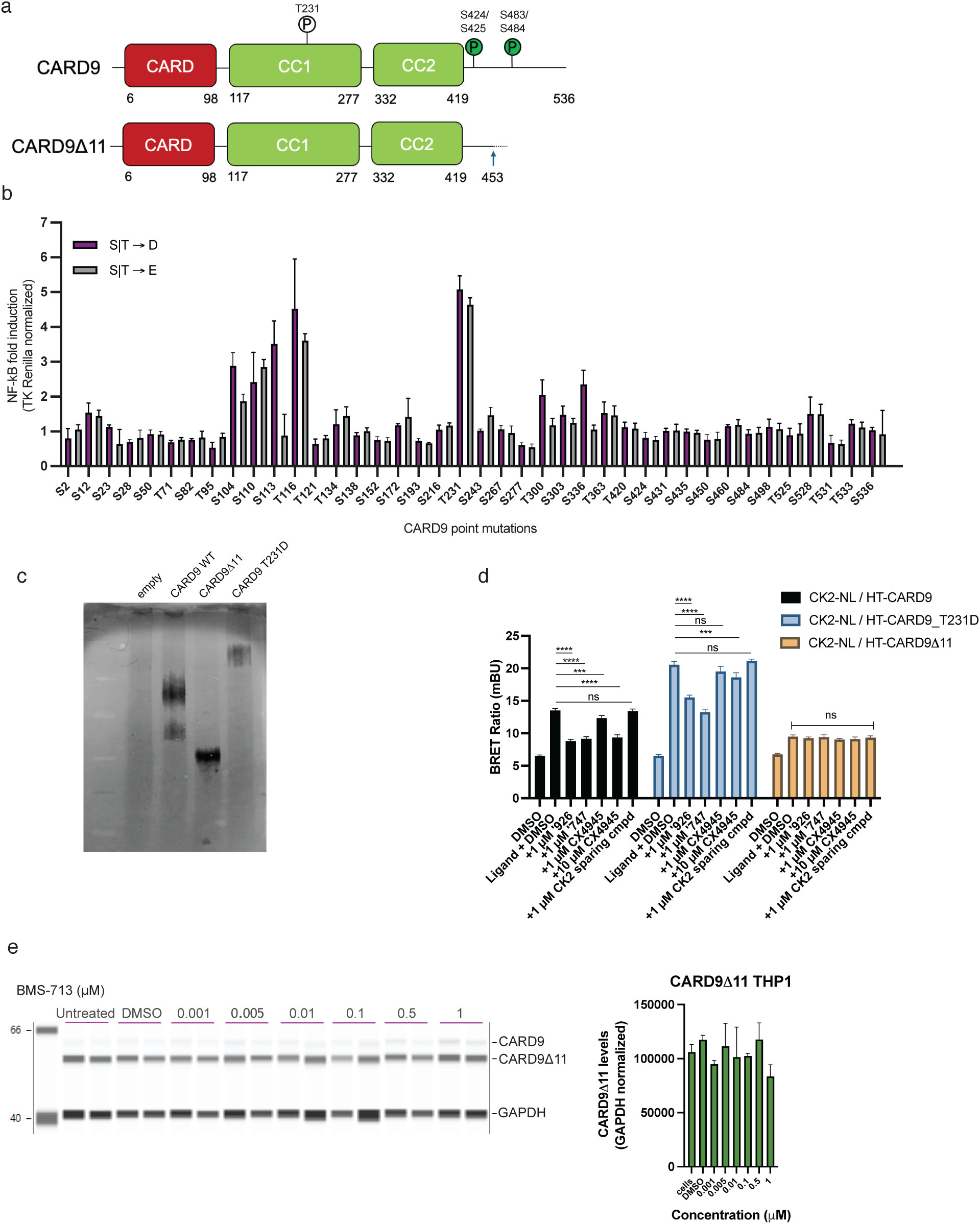
CARD9Δ11 does not bind to CK2 and is protected from depletion. **(a)** CARD9 and CARD9Δ11 domain architecture as annotated in Uniprot with PKCδ (T231) and CK2 (S424/S425 and S483/S484) phosphorylation sites depicted. **(b)** Phosphomimetic scanning (S|T → D|E) of CARD9 protein (n = 3) identifies gain of function (GoF) mutations. **(c)** CARD9 T231D GoF exhibits larger oligomeric species by native gel. **(d)** CK2 inhibitors reduce BRET interaction between CK2-NL and HT-CARD9 or HT-CARD9 T231D but not HT-CARD9Δ11 (4-hour incubation). Data points are reported as mean ± SD (n = 4). Statistical significance determined by one-way ANOVA with adjusted p value depicted (*-*p* ≤ 0.05, **-*p* ≤ 0.01, ***-*p* ≤ 0.001, ****-*p* ≤ 0.0001). **(e)** Jess Western blot of CK2i titration of endogenously edited THP-1 CARD9Δ11 cell line, which does not exhibit CK2 inhibition-driven depletion of CARD9Δ11 (overnight incubation).

Phosphomimetic mutations to CK2 phosphorylation sites, S424 and S484, did not result in any significant effects on NF-κB induction. However, we found that mutagenesis to the predicted bridge between the CARD domain and first coiled-coil domain (S104, S110, S113, and T116) as well as T231 conferred significant induction of NF-κB (Fig. 5b). The T231 amino acid is the precise residue phosphorated by PKCδ to activate CARD9^2^, making the T231D or T231E mutation an appealing way of genetically encoding CARD9 gain-of-function (GoF) without upstream stimulation of the pathway. We observed by Western blot that the T231D and T231E mutations conferred less speciation of CARD9 and a relative preference for higher order oligomers compared to WT CARD9 (Supp. Fig. 7). The preference for higher order oligomers by CARD9 T231D was confirmed by native gel (Fig. 5c) and supported its utility as a CARD9 GoF construct based on its oligomerization-driven mechanism of action.

### CARD9Δ11 is protected from CK2-dependent CARD9 depletion

To further examine the mechanism behind CARD9-CK2, we leveraged the CARD9 GoF construct (T231D) and the CARD9 LoF construct (CARD9Δ11) in our nanoBRET assay. CARD9 WT exhibited loss of BRET with CK2 when incubated with our CK2 inhibitor lead series and CX4945 but not with a CK2 sparing compound (Fig. 5d, Fig. 4e). A similar dependence was found with CARD9 T231D and CK2, yet the magnitude of BRET between CK2 and CARD9 T231D was higher than with CARD9 WT, suggesting a stronger interaction between the two. Notably, BRET between CK2 and CARD9Δ11 was significantly attenuated compared to CARD9 WT and exhibited no change when incubated with CK2 inhibitors, suggesting that CK2 cannot bind or does not bind well to the CARD9Δ11 protein.

The CARD9Δ11 allele is truncated at the C terminus of CARD9, leaving only the first 453 of 536 amino acids conserved with the WT allele. Since the CARD9Δ11 truncation is localized to the region where CK2 phosphorylates CARD9 (S424, S425, S483, and S484), we hypothesized CK2 may not act on CARD9Δ11 as a substrate, and therefore CK2 inhibition may not affect CARD9Δ11 protein levels. To test this, we endogenously edited the intronic region of CARD9 using CRISPR/cas9 in a THP-1 cell line to induce the expression of CARD9Δ11. When we incubated the CARD9Δ11 THP-1 cell line with BMS-713, we discovered that the CARD9Δ11 protein was not depleted as seen in the WT THP-1 cell line (Fig. 5e, Fig. 2c), suggesting that CK2 may not stabilize the CARD9Δ11 protein in carriers of that allele.

### CK2 inhibition depletes CARD9 in murine bone marrow-derived dendritic cells

To ultimately assess whether CK2 inhibition causes CARD9 depletion in vivo, we first needed to evaluate whether the CARD9-CK2 mechanism is conserved in mice. The CARD9-CK2 phosphorylation sites, S424 (mS424), S425 (mS425), S483 (mS483), and S484 (mS484), are conserved in the CARD9 murine sequence, and we observed potent CARD9 depletion in bone marrow-derived dendritic cells (BMDCs) when incubated with our CK2 inhibitors (Fig. 6a). The DC_50_ and Y_min_ values tracked similarly to those observed in human dendritic cells with our lead series (DC_50_ <50 nM and Y_min_ <50%). CX4945 exhibited similar micromolar potency (∼4 μM) as observed in THP-1s (∼4 μM) and human primary dendritic cells (∼2 μM). As previously observed in human dendritic cells, the CK1 inhibitor, BMS-321, did not deplete CARD9 in BMDCs, reinforcing that the CARD9 depletion mechanism by CK1 is unique to immortalized cell lines.

**Fig. 6.**
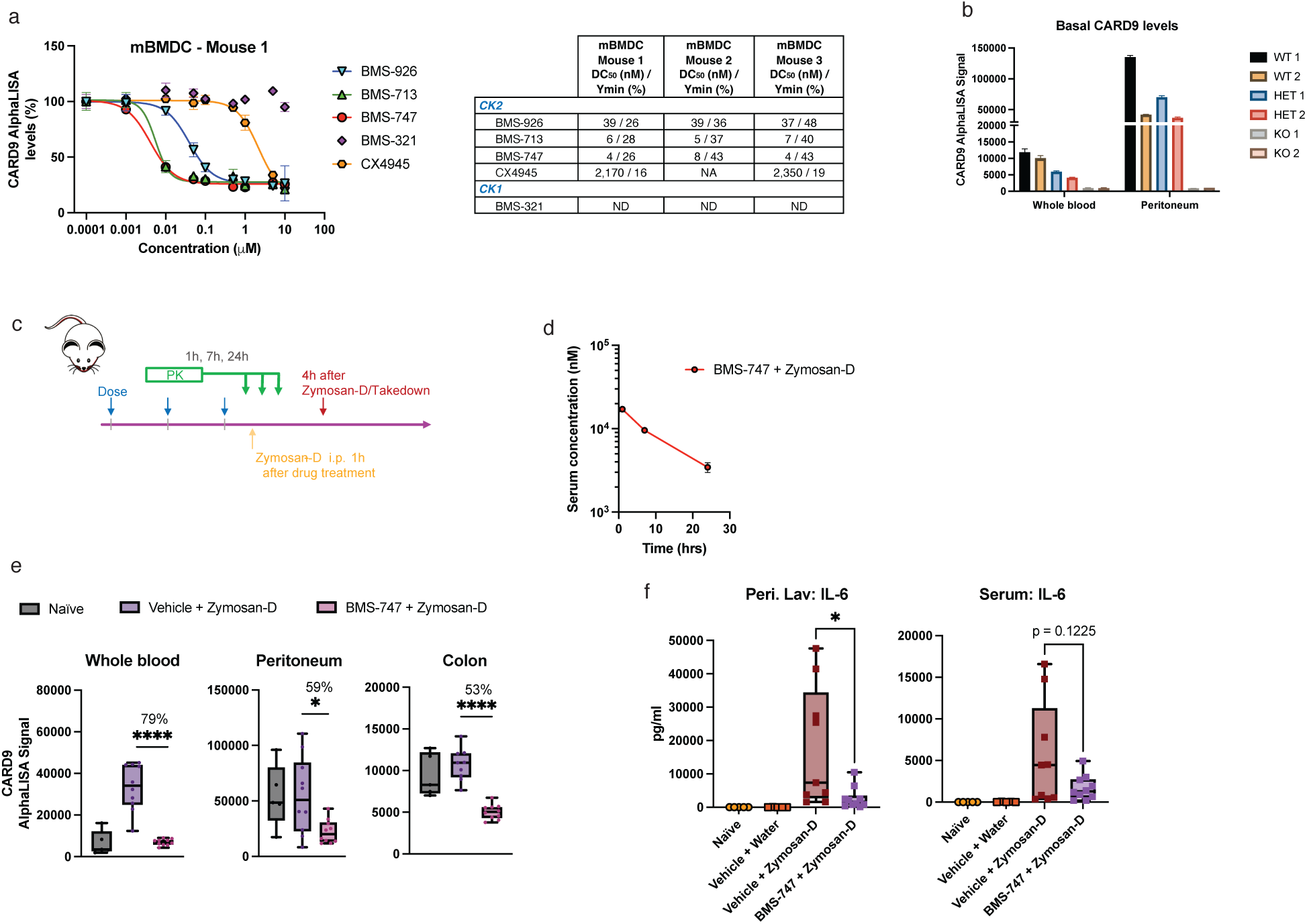
CK2 inhibition drives CARD9 depletion *in vivo*. **(a)** CK2 inhibition leads to CARD9 depletion in bone marrow-derived dendritic cells (BMDCs) as measured by CARD9 AlphaLISA (overnight incubation). Data represented as mean with SD in graph. **(b)** Relative quantitation of CARD9 in various murine compartments of wild type (WT), heterozygous CARD^+/−^, and CARD9^−/−^ KO mice by CARD9 AlphaLISA. **(c)** Study design to evaluate *in vivo* CARD9 depletion driven by CK2 inhibition. **(d)** Serum exposure of BMS-747 1, 7, and 24 hr after the penultimate dose of BMS-747. measured by LC-MS. **(e)** AlphaLISA CARD9 levels in the whole blood, peritoneal lavage cells, and colon from naïve (n = 5), vehicle + zymosan-D (n = 10) and BMS-747 + zymosan-D (n = 10) treated mice. Statistical significance determined by one-way ANOVA with adjusted p value depicted (*-*p* ≤ 0.05, **-*p* ≤ 0.01, ***-*p* ≤ 0.001, ****-*p* ≤ 0.0001). **(f)** Luminex IL-6 levels in the peritoneal lavage and serum of mice at termination. Statistical significance determined by one-way ANOVA with adjusted p value depicted (*-*p* ≤ 0.05, **-*p* ≤ 0.01, ***-*p* ≤ 0.001, ****-*p* ≤ 0.0001).

### CK2 inhibitors deplete CARD9 *in vivo*

To examine whether CK2 inhibition impacts CARD9 protein homeostasis *in vivo*, we needed to establish reliable CARD9 detection *ex-vivo*. We found that AlphaLISA could reliably detect CARD9 *ex-vivo* within murine compartments such as: whole blood, colon, and peritoneum (Fig. 6b, Fig. 6e). The highest abundance of CARD9 was observed in the peritoneal cavity, potentially due to the high myeloid cell population.

BMS-747 demonstrated excellent PK properties that covered the apparent IC_50_ over 24 hours (Fig. 6c, 6d) and was subsequently chosen for *in vivo* study. To evaluate the effect of CK2 inhibition on CARD9 levels, BMS-747 was administered orally (po) at 10 mg/kg for 3 days QD followed by an injection of zymosan-D (50 mg/kg) intraperitoneally (ip) on day 3 and euthanization 4 hours later (Fig. 6c). Significant CARD9 depletion was observed in the whole blood, colon and peritoneum compartments (Fig. 6d). CARD9 levels rose significantly in mouse whole blood with zymosan-D administration, but this effect was not seen in other compartments. Notably, IL-6 levels were also significantly attenuated in the peritoneal lavage fluid and trended lower within mouse serum (*p* = 0.1225). Taken together, these results indicate that the CARD9-CK2 mechanism is biologically conserved between mouse and human and offers an unconventional approach for engaging a therapeutically intractable target.

## Discussion

Here we discover that CARD9 is a substrate for CK2 and directly influences CARD9 protein homeostasis (Fig. 7) with the promise that this mechanism could be leveraged to treat autoimmune disease. It is noteworthy to mention that we found that CK2 inhibition does not impact the protein homeostasis of CARD9Δ11, which does not bind to CK2 and partially lacks CK2 phosphorylation sites. It has previously been reported that CARD9Δ11 acts as a dominant negative LoF construct, which leads to protection within autoimmunity.^4^ Our data suggests an additional mechanism that the CARD9Δ11 allele uses to confer protection in which the protein is destabilized, leading to apparent lower expression and reduced cytokine induction.

**Fig. 7.**
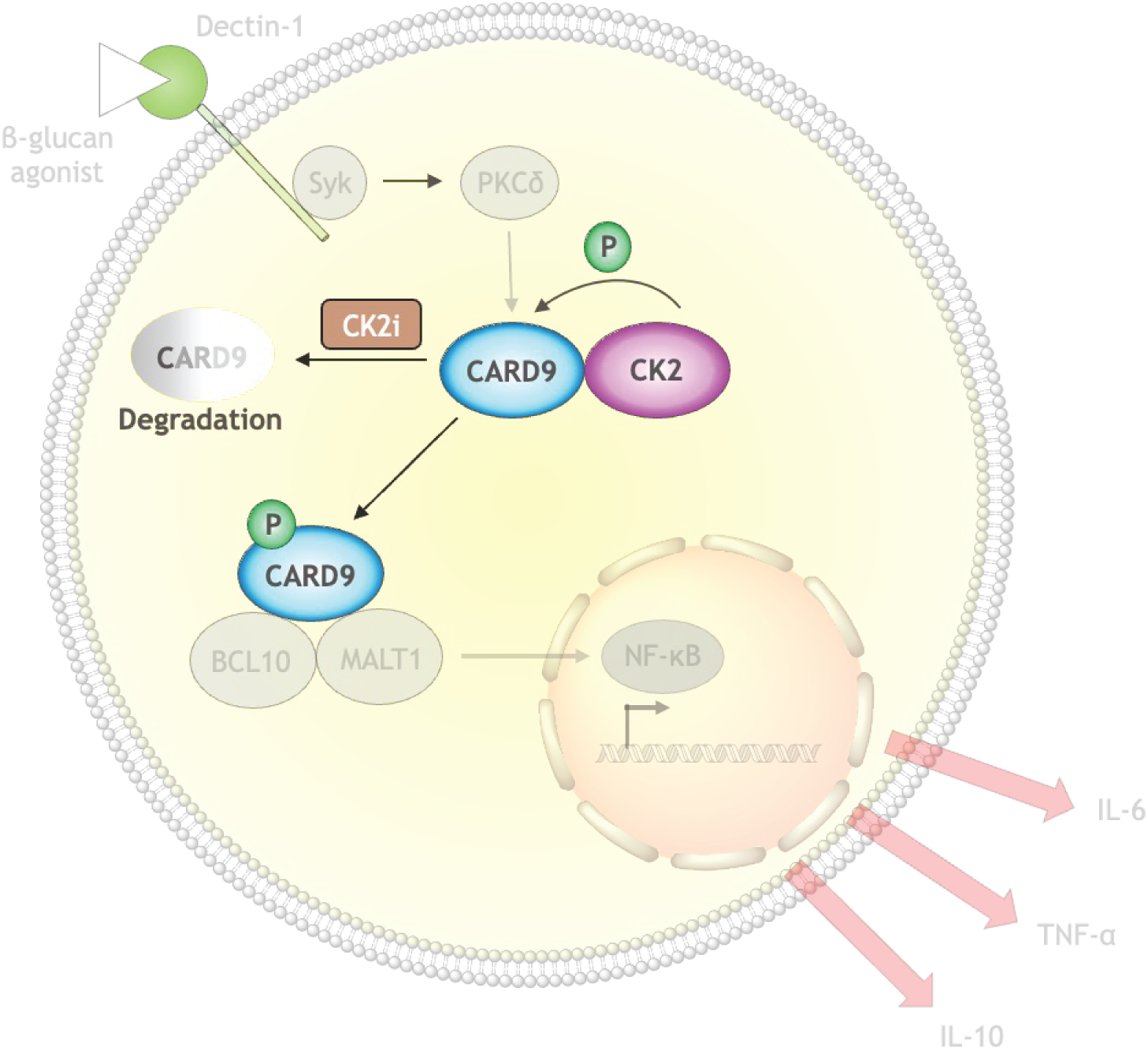
CARD9 pathway depiction of CARD9-CK2 interaction and downstream effects.

CK2 is a complex target capable of phosphorylating hundreds of substrates and influencing a multitude of signaling pathways. This work also expands the scope of CK2 phosphorylation substrates and adds to a growing list of mechanisms of action influenced by CK2 that modulate immune cell activity. It has previously been shown that CX4945 promotes T_reg_ differentiation through simultaneous pSTAT3 inhibition and pSTAT5 activation in a murine trinitrobenzene sulfonic acid (TNBS)–induced colitis model.^28^ Similar findings were reported in experimental autoimmune encephalomyelitis (EAE) murine models where CK2 genetic ablation or inhibition promoted T_reg_ development while preventing Th17 development.^29^ In addition, CK2α has been implicated in CD4^+^ T cell proliferation by regulating NFAT2, which has been shown to contribute to the pathogenicity of colitis.^30^ Therefore, the previously unappreciated link between CARD9 and CK2 deepens our understanding of how CK2 inhibition impacts fundamental immunology as well as within the context of disease.

The impact of CK2-dependent phosphorylation on protein homeostasis is not without precedent. It has been shown that CK2 binds and phosphorylates PERIOD2 (PER2) S53, a key protein for maintaining the circadian clock.^31^ S53 phosphorylation of PER2 can act independently or in concert with CK1ε-dependent phosphorylation of PER2, leading to PER2 protein degradation.^31^ Other CK# isoforms have also been reported to stabilize or destabilize its binding partners by phosphorylation. Bidère et al. discovered a protein-protein interaction between CK1α and CARD11 and stated, “…our preliminary data indicate that CK1α might contribute to CARMA1 [CARD11] degradation”.^32^ Dushukyan and colleagues demonstrated that CK1ε activates protein phosphatase 5 (PP5), and in the absence of phosphorylated PP5, CK1ε inhibition promotes VHL-mediated ubiquitination and subsequent degradation of PP5.^33^ Finally, a global comparative analysis of phosphorylated and non-phosphorylated structures in the Protein Data Bank (PDB) indicates that protein phosphorylation generally introduces small conformational changes that are stabilizing.^34^ Kinase inhibitors that destabilize kinases were long thought to operate through chaperone deprivation.^35^ However, Scholes and colleagues have recently illuminated that some kinases can be degraded irrespective of the chaperone status and without direct binders.^36^ Taken together, these endogenous mechanisms to stabilize proteins via phosphorylation can be leveraged to deplete proteins of interests by small molecule-mediated inhibitions.

CK2 inhibition has been implicated as a therapeutic target for a wide spectrum of diseases including neurodegenerative, COVID-19 infection, cancer, cardiovascular, and autoimmune.^37^ In the clinic CK2 inhibitors have been evaluated for cancer and COVID-19 with limited adverse effects. While there is a potential for leveraging CK2 pharmacological inhibition to modulate CARD9-dependent pathology, the promiscuity of CK2 makes the identification of a single causal mechanism of action difficult. This creates a need to interpret the ensemble of effects from CK2 inhibition that overlap with further studies to elucidate the exact mechanism of CK2-dependent CARD9 depletion. Our work also invites biological questions on the connection between CARD9 and CK#. Given the roles of CK1 and CK2 in mammalian circadian rhythms, we hypothesize that regulation of CARD9 expression by CK# is a single component of a larger CK#-dependent influence on the circadian immune system.^38^ While this area is poorly understood, this could confer a periodicity to CARD9 regulation and serve as a rich area for further study.

## SUPPLEMENTAL FIGURES

**Supplemental Fig. 1.**
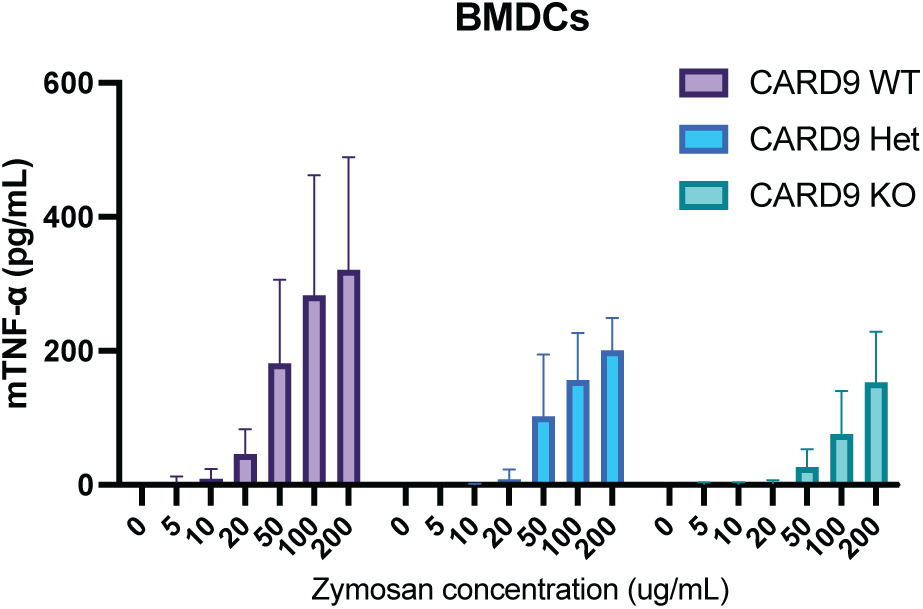
TNF-α in BMDCs by zymosan stimulation is dependent on CARD9 levels (n = 3). Data represented as mean with SD in graph (overnight incubation).

**Supplemental Fig. 2.**
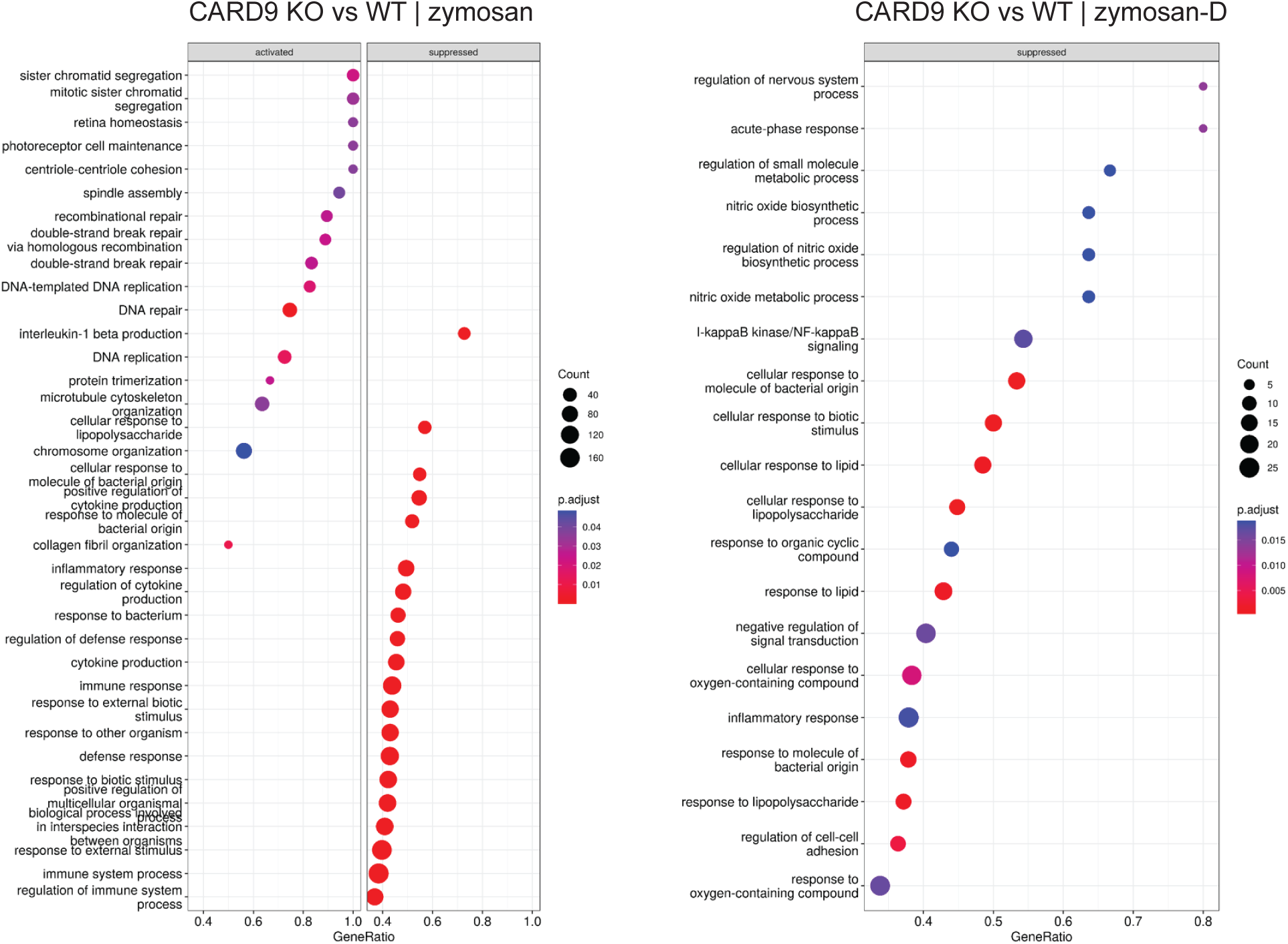
Gene ontology analysis of WT vs CARD9 KO murine neutrophils treated with zymosan or zymosan-D.

**Supplemental Fig. 3.**
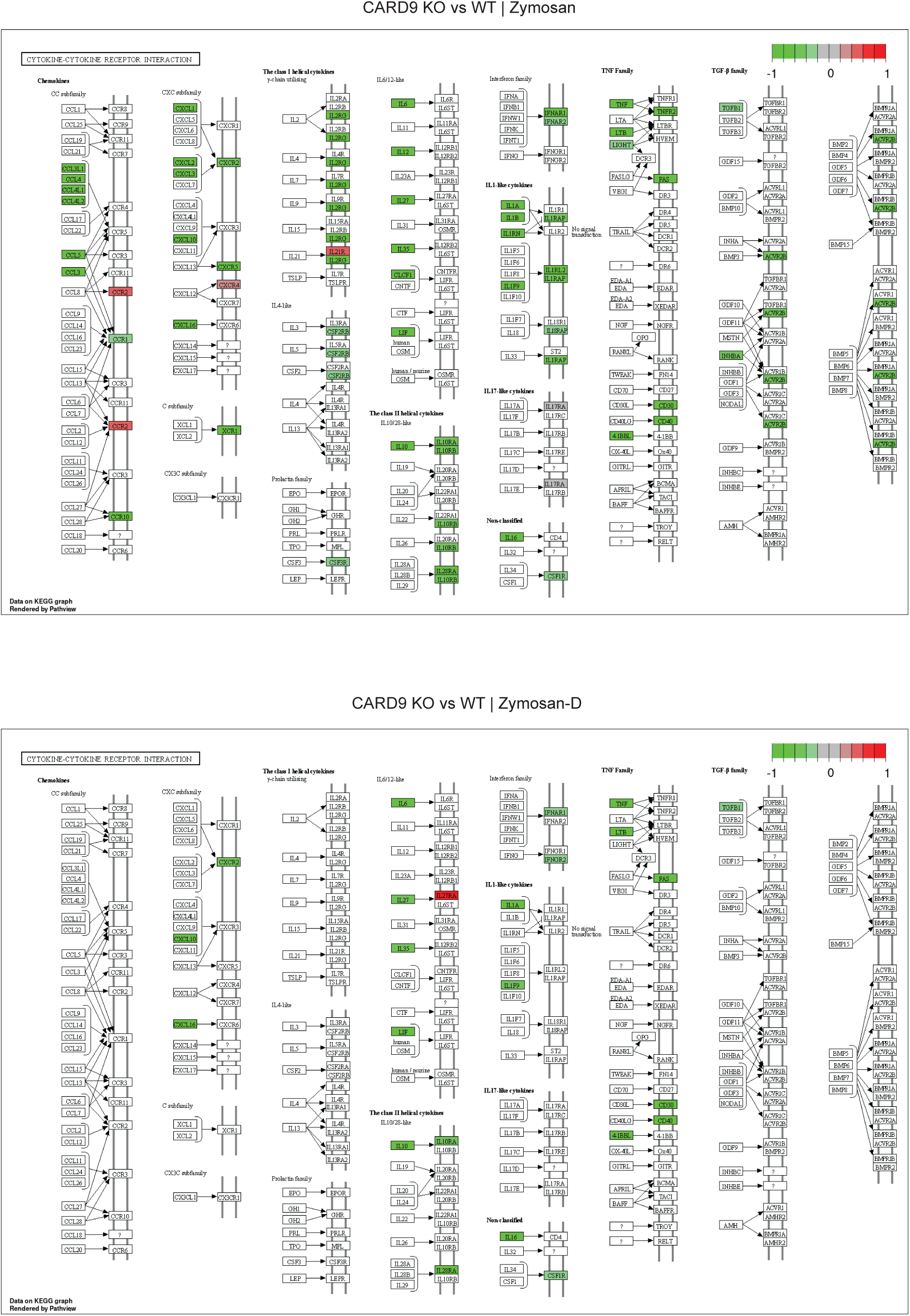
Cytokine pathway analysis comparing WT vs CARD9 KO murine neutrophils treated with zymosan or zymosan-D.

**Supplemental Fig. 4.**
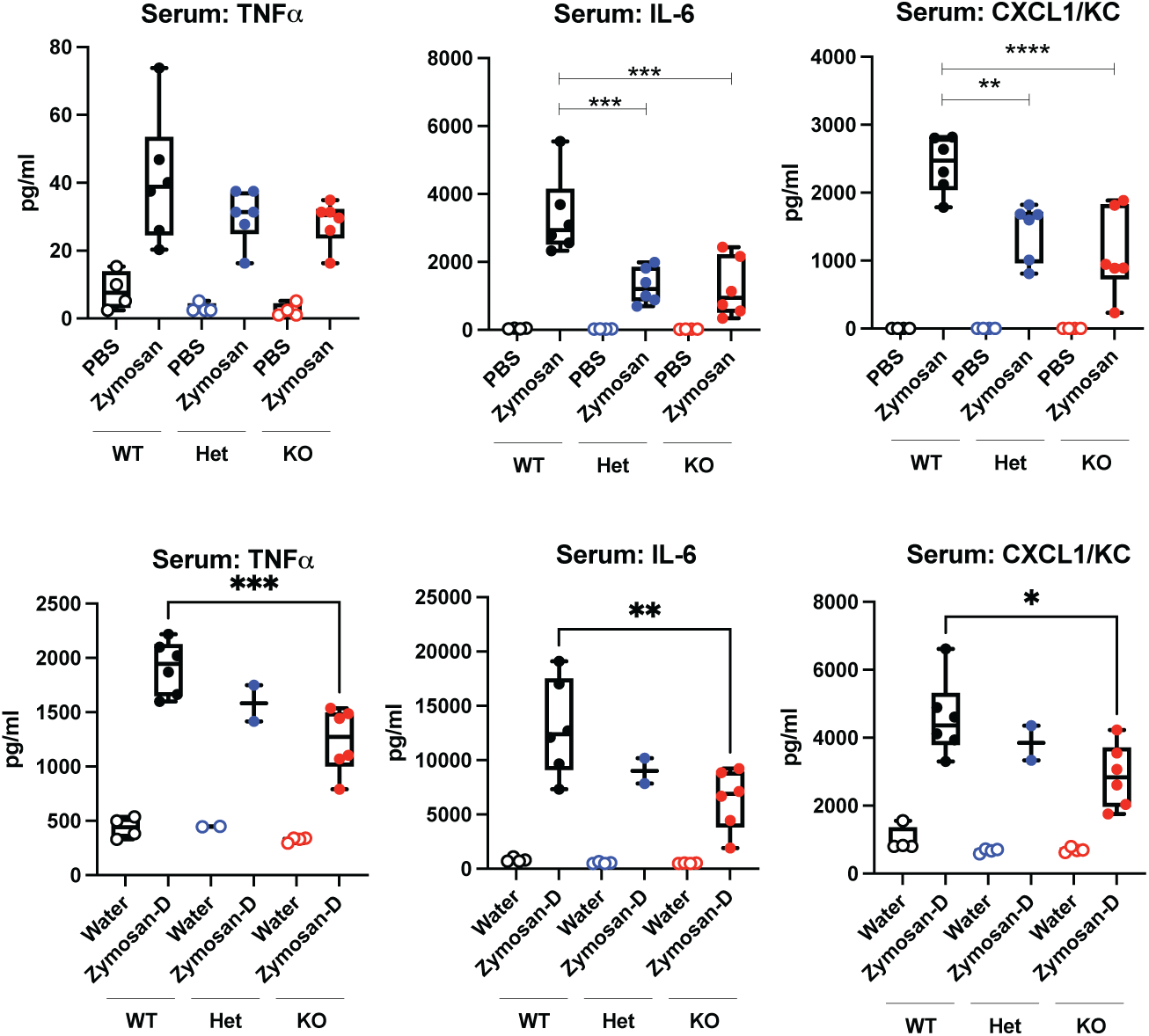
CARD9 loss shows attenuation of pro-inflammatory cytokine induction in zymosan mouse model. Luminex levels of TNF-α, IL-6, and CXCL1/KC in the serum of either PBS (n = 4), zymosan (50 mg/kg, n = 6), water (n = 4), or zymosan-D (50 mg/kg, n = 6)-treated WT, CARD9^+/−^ heterozygous or CARD9^−/−^ KO mice. Statistical significance determined by one-way ANOVA with adjusted p value depicted (*-*p* ≤ 0.05, **-*p* ≤ 0.01, ***-*p* ≤ 0.001, ****-*p* ≤ 0.0001).

**Supplemental Fig. 5.**
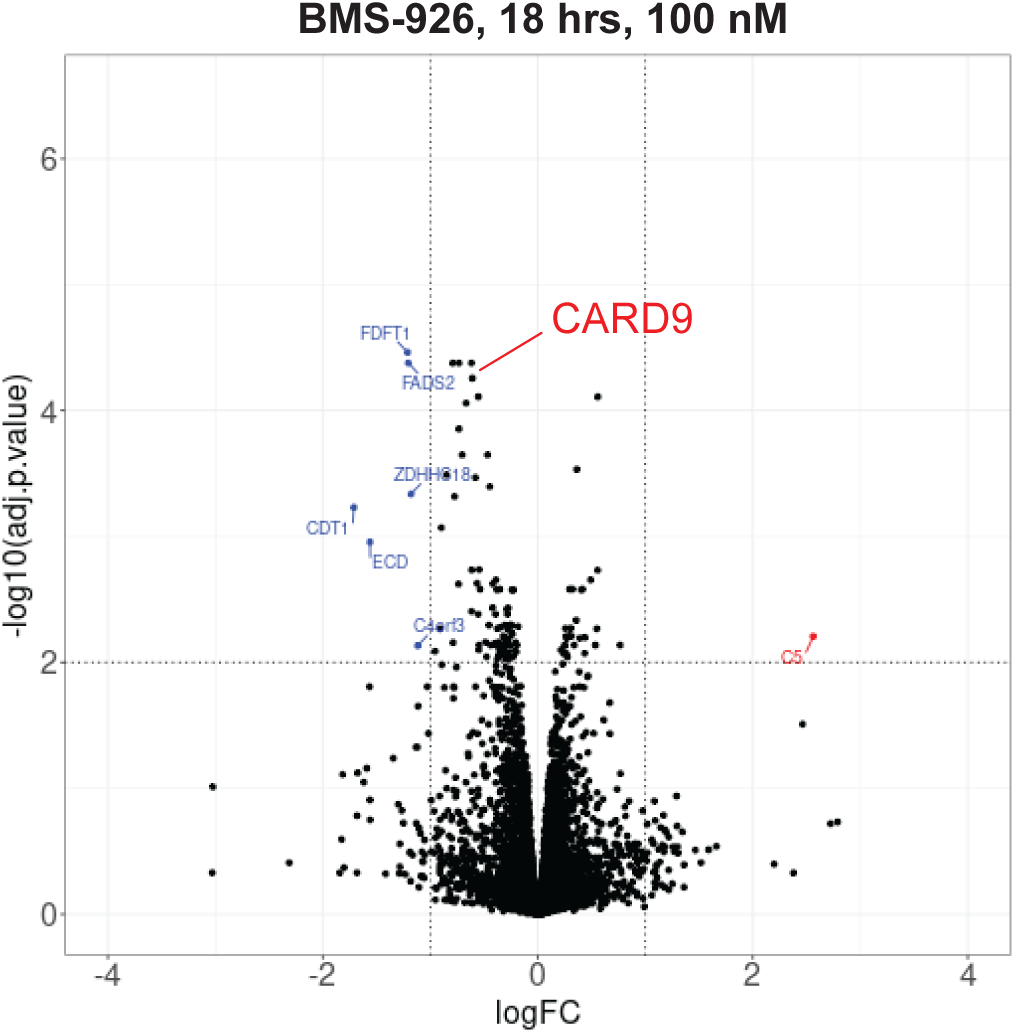
Global proteomics of CK2i treatment in THP-1 cell line.

**Supplemental Fig. 6.**
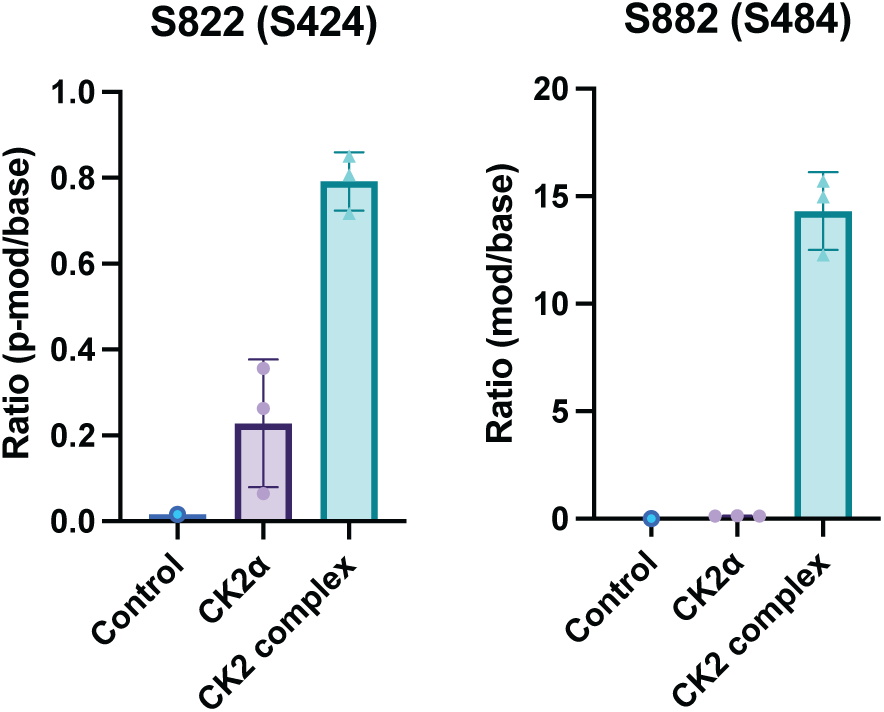
Identification of CARD9 phosphorylation sites by CK2. Abundance ratio of the monophosphorylated peptide over unmodified peptide from MBP-CARD9 containing the indicated serine residue.

**Supplemental Fig. 7.**
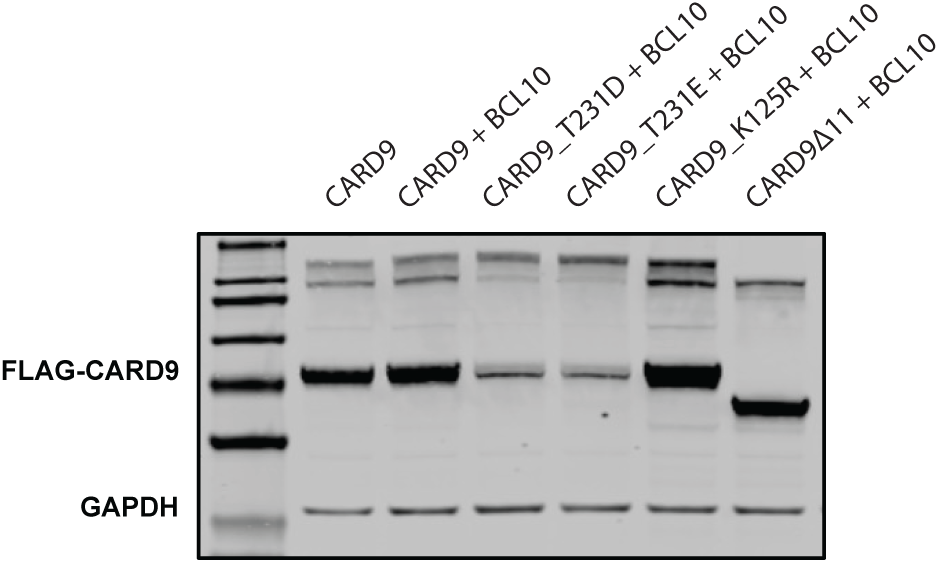
Western blot shows higher order speciation of CARD9 oligomers. HEK293T cell line transiently transfected with indicated constructs and blotted for FLAG tag. All CARD9 constructs harbor N-terminal FLAG tag.

## ACKNOWLEDGMENTS

The authors thank Ping Chen, Radhika Goenka and Jenny Xie for *in vivo* discussions and study designs. We thank Pathik Rakesh Desai for spearheading the creation of the CARD9 AlphaLISA kits.

## AUTHOR CONTRIBUTIONS

## Methods and Materials

### Cell Culture and Reagents

All cells were grown in a 37 °C, 5% CO_2_, humidified environment, in media containing 1% penicillin/streptomycin (Pen/Strep). Cell lines THP-1 (ATCC) and SNU-C1 cells (ATCC) were grown in RPMI with 10% heat-inactivated fetal bovine serum (HI-FBS), L929 (ATCC) in EMEM with 10% HI-FBS, HEK293T (ATCC) and Lenti-HEK (Takara) in DMEM with 10% HI-FBS. Endogenously edited THP-1, including HiBiT-CARD9, CARD9 T231D, CARD9Δ11, were generated at Synthego.

### Mice

All protocols and studies were approved by the Bristol Myers Squibb IACUC committee. CARD9^WT^, CARD9^HET^ and CARD9^KO^ mice were generated and bred at Jackson Laboratory. C57BL/6 mice were purchased from Charles River Laboratory. Female mice were used in all studies at greater than 8 weeks of age. Mice were allowed to acclimate for at least 72 hrs before the study. Identical enrichment was provided for each cage: InnoWheel and Bed’r nest; along with standard corn cob bedding and rodent chow. CARD9^WT^, CARD9^HET^ and CARD9^KO^ mice were ear notched using Stoelting™ Ear Punch (Fisher Scientific) and genotyping was verified by Transnetyx. For all in vivo studies, mice were monitored daily for health checks and weighed daily.

### Mouse Peritonitis Model

In the peritonitis study, C57BL/6 mice were administered 50 mg/kg of zymosan (Invitrogen, tlrl-zyn)) or Zymosan-D (Invitrogen, tlrl-zynd) or its vehicle intraperitoneally and harvested 4 hrs later. Mice were euthanized using CO_2_ and cervical dislocation. Upon confirmation of death, cardiac puncture was carried out to collect whole blood and placed in either lithium heparin coated tubes or serum separator tubes. Serum was isolated using serum separator tubes. Collected blood was stored on ice for a maximum of 30 min prior to centrifugation at 8000 rpm for 8 min and separation. Peritoneal lavage was performed with one injection of 5 mL of EDTA-PBS (Teknova), and once collected, tubes were spun at 1500 rpm for 5 min and the supernatant and pellets were separated and stored frozen until analysis. To analyze mediators in peritoneal lavage and serum, Luminex 26-plex or IL-6 single-plex kits (ProcartaPlex™ Mouse Cytokine & Chemokine Convenience Panel 1, 26-plex, and IL-6 Mouse ProcartaPlex™ Simplex Kit, Thermofisher) were used.

To evaluate the effect of BMS-747 in the peritonitis model, BMS-747 (10 mg/kg) or its vehicle (4 mL/kg; 5% Dimethyl Acetamide; 9.5% Ethanol; 76% PEG-400; 9.5% Vitamin E TPGS) were delivered orally once a day for 3 days. Serum was collected via submental bleeds at 1, 7 and 24 hrs after the penultimate dose of BMS-747 and the serum concentration of BMS-747 was quantified in serum by LC-MS Sciex (6500+). One hour after the last dose of BMS-747 or vehicle, mice received zymosan-D challenge at 50 mg/kg, and euthanized 4 hrs later. In addition to whole blood, peritoneal lavage collected as described above, colons were excised from the cecum to the rectum, flushed with PBS to remove fecal matter and stored frozen until analysis.

Data are expressed as mean (±SEM). Differences between means were tested for statistical significance on GraphPad Prism using one-way ANOVA test for multiple comparisons between all groups (Tukey’s test for multiple comparisons). From such comparisons, differences yielding *P* value less than 0.05 were judged to be significant.

### Mouse Primary Cell Culture

Mouse bone marrow derived dendritic cells (BMDC) were obtained from extracted mouse femurs. Once femurs were isolated, the bone marrow was extracted using mortar and pestle and filtered through a 70 μm filter to isolate the cells from the debris. The cells were cultured in RPMI with 10% HI-FBS and 1x Pen/Strep. To differentiate the cells into BMDCs, they were cultured in media supplemented with 10% L929 granulocyte colony-stimulating factor enriched media and 20 ng/mL of purified mouse IL-4 (Gemini Bio). L929 enriched media was obtained from culturing L929 cells until they were confluent and resting them for 10 days before taking off the enriched media and spin purifying it. DCs differentiated for 7-10 days before being used for experiments. To isolate the DC adherent population from potential macrophages, Versene solution (Gibco) was applied to PBS washed cells and allowed to sit at room temperature for 5 min before being vigorously pipetted off the plate, leaving behind any macrophages. Mouse neutrophils were obtained from mouse femur bones where bone marrow was extracted as described above. The neutrophils were then isolated from the marrow cells using EasySep™ Mouse Neutrophil Enrichment Kit (Stemcell, 19762) following the manufacturer’s instructions.

### Cytokine Analysis of Zymosan or Zymosan-D Treated BMDCs and Neutrophils

50-100k cells were plated in a 96-well plate, and allowed to rest for 24 hrs prior to stimulation, and then treated with either zymosan or zymosan-D for indicated time and concentrations. Supernatants were collected by centrifuging cells at 800 x g for 5 mins. Subsequent TNFa and IL-6 AlphaLISAs (Revvity) were conducted according to manufacturer’s specifications for cytokine detection.

### RNA-seq Analysis of Zymosan and Zymosan-D Treated Mouse Neutrophils

1e^6^ mouse neutrophils from CARD9^WT^ and CARD9^KO^ mice were treated with 100 μg/mL zymosan or zymosan-D for 2 hrs as described above, and cell pellets were harvested. RNA isolation and subsequent RNA-seq analysis was performed at Azenta.

Out of 27,223 genes detected in at least one sample in the RNA-seq study, 13,941 genes were removed because they were lowly expressed (less than 5 counts in at least 10% of the samples). The remaining 13,282 genes were carried forward for downstream analysis. Principal component analysis (PCA) was performed using the *prcomp()* function from *stats* packaged in R.

Prior to differential expression analysis (DEA), unknown confounding effects within the data were identified using Surrogate Variable Analysis (SVA).^1^ The expression data were modeled using the formula ∼ *Treatment* ∗ *Genotype*, where *Treatment* is a categorical variable with the levels “None”, “Zymosan”, and “Zymosan-D”, and *Genotype* is a categorical variable with the levels “WT”, “CARD9 Heterozygous KO”, and “CARD9 Homozygous KO” against a null model (∼ 1). *svaseq()* function from the *sva* package identified 7 surrogate variables that capture confounding effects outside the defined model parameters. None of these variables correlated with the *Treatment* or *Genotype* variables and therefore they were all included in the final model as additional confounding variables. DEA was performed on log2 normalized data using the *DEseq2* package in R based on the final model of ∼ *Treatment* ∗ *Genotype* + *Surrogate Variables* [1 − 7]. P-values were corrected for multiple testing using false discovery rate (FDR) method via *qvalue()* function. Significant genes were selected at a q-value threshold of 0.01 and a log2 fold change cutoff of 1 for each contrast. Volcano plots for the differentially expressed genes were generated using the *EnhancedVolcano* package in R. Expression heatmaps were generated using the *pheatmap()* function from the *pheatmap* package. Venn diagrams of overlapping gene targets between different contrasts of interest were generated using the *ggvenn* package in R.

GSEA was performed for each contrast separately, on differentially expressed genes using the *gseGO()* function from the *clusterProfiler* R package. “org.Mm.eg.db” was used for the gene set annotations, “fdr” was used as multiple testing correction method, and 0.05 was the cutoff for significant enrichment. KEGG pathway enrichment analysis was performed using *gseKEGG()* function with the same cutoff parameters. Individual significant pathways were visualized using *pathview* package.

### HiBiT Screening

In a 1536-well plate, compounds were transferred via Echo (labcyte) to make final concentration of 5 μM, and suspension of THP-1 HiBiT-CARD9 cells in the assay buffer (phenol-red free RPMI, 2% HI-FBS, 1x Pen/Strep) were added to achieve 2500 cells per well (6 μL at 0.833 e^6^/mL). After 22 hr incubation at 37 °C, CellTiter-Blue (Promega, G8080) was added to the cells. Fluorescence was measured after 2 hr incubation at 37 °C at 560/590 nm excitation/emission on EnVision (Revvity), after which Nano-Glo Hibit Lytic reagent (Promega, N3040) was added to read luminescence on EnVision.

### AlphaLISA Screening

CARD9 AlphaLISA kits were custom made by Revvity, and one kit was utilized to detect human CARD9 and another for mouse CARD9. Unless noted otherwise, CARD9 levels in lysates or tissue homogenates were detected by following manufacturer’s instructions.

In a 384-well plate, compounds were transferred via Echo to achieve desired concentrations. THP-1 cells were added in the assay buffer (phenol-red free RPMI, 2% HI-FBS, 1x Pen Strep) to yield 10k cells/well (20 μL at 0.5 e^6^/mL). Following overnight incubation, 5 μL of 5x AlphaLISA Lysis (Revvity, AL003F) buffer was added to the cells, and allowed to lyse at room temperature for 30 min with shaking. 2 μL of resultant lysate were transferred to Proxiplate (Revvity), and then incubated with 4 μL of the mixture of the biotin-antibody and the acceptor beads at room temperature for 1 hr. 2 μL of the donor beads were then added and the plates were incubated in the dark overnight then read on EnVision at 615 nm. In parallel, cell viability was assessed with CellTiter-Glo (Promega, G7573)) per manufacturer’s protocol.

### Global Proteomics

THP-1 cells were seeded in a 96-well plate and treated with the compound for the indicated time. Tryptic lysates were prepared for proteomic data acquisition using the PreOmics iST-BCT 96 sample kit. Protein digests were subjected to data collection using an LC-MSMS system (Evosep, #EV-1000) connected online to a mass spectrometer (Bruker, timsTOF pro2) in Data Independent Parallel Accumulation Serial Elution Fragmentation mode (DIA-PASEF). Peptide identification and relative quantification were performed using DIA-NN v1.8.1^2^ with an experiment specific library generated from THP-1 cells. Precursor data obtained from DIA-NN was subjected for imputation with minimal detection imputation (MinDet) via the imputeLCMD package,^3^ setting a quantile threshold at 0.01. Differential expression analysis was performed on the imputed data with using the limma package.^4^

### CK2 Functional Assay via SNU-C1 Cell Proliferation

SNU-C1 cells were added to a 384 well plate in the assay buffer (RPMI 2% HI-FBS 1x Pen Strep) at 1000 cells/well (40 μL at 25k/mL) and incubated overnight. Compound were added to cells via Echo, and cell plates were incubated for 72 hrs. Four microliter of MTS reagent (Promega, G5421) was added and the plates were read on EnVision at 490 nm. Campothecin was used as positive control.

### Arrayed CRISPR Screen

THP-1 HiBiT-CARD9 + Cas9 cells were generated by introducing Cas9 and Blasticidin resistance expression with lentivector pLX_311-Cas9 (Broad Functional Genomics Consortium). THP-1 HiBiT-CARD9 + Cas9 cells were grown in RPMI with 10% HI-FBS, 1x Pen/Strep, 2-mercaptoethanol, and 3 µg/mL Blasticidin. Arrayed lentivirus containing sgRNA constructs in lentivectors pXPR_003 or pXPR_050 were purchased from the Broad Functional Genomics Consortium. 20k THP-1 HiBiT-CARD9 +Cas9 cells were plated in a round bottom 96-well plate (Corning) and transduced with arrayed virus (Broad Functional Genomics Consortium) in media supplemented with 4 µg/mL polybrene (Millipore Sigma) by centrifugation at 1200 x g for 30 min. After overnight incubation, media was removed and replaced with fresh media. The following day cells were selected with puromycin. Seven days after transduction, media was changed and 25 µL of cells were rearranged in quadruplicate into two 384-well plates. The remainder of the culture was passaged for analysis on day 14 to observe cells in log phase growth. Nano-Glo HiBiT and CellTiter-Glo assays were performed by addition of 25 µL Nano-Glo HiBiT lytic detection mix (Promega) or CellTiter-Glo reagent (Promega) to each well, mixed for 5 min, incubated for 10 min, and measured on an EnVision plate reader.

### Human Primary Cell Culture

Human monocytes were isolated from fresh whole blood of healthy donors (Bristol Myers Squibb internal donor program). After whole blood was collected in EDTA tubes and monocytes were isolated using EasySep™ Direct Human Monocyte Isolation Kit (Stemcell, 19669) following the manufacturer’s instructions. Dendritic cells were obtained by culturing isolated monocytes in RPMI with 10% HI-FBS, 1x Pen/Strep with 50 ng/mL GM-CSF (Gemini Bio) and 20 ng/mL of purified human IL-4 (Gemini Bio). DCs differentiated for 7-10 days before being used for experiments. To isolate the DC adherent population from potential macrophages, Versene solution (Gibco) was applied to PBS washed cells and allowed to sit at room temperature for 5 min before being vigorously pipetted off the plate leaving behind any macrophages.

### CARD9 Quantification in hDC or mBMDC via AlphaLISA

50-100k cells were plated in a 96-well plate and allowed to rest. Using a HP D300 Tecan liquid dispenser, compounds were added to the cell suspension. After overnight incubation, cells were harvested by centrifugation at 500 x g for 5 min, and cells were lysed in AlphaLISA buffer and allowed to lyse for 30 min at 4 °C. Human AlphaLISA kit for hCDs and mouse AlphaLISA kit for mBMDCs were performed using manufacturer’s instructions.

### CARD9 Quantification in hDC or THP1 CARD9 Δ11 via JESS Western Blot

50-100k cells were plated in a 96-well plate and allowed to rest. Using a HP D300 Tecan liquid dispenser, compounds were added to the cell suspension. After overnight incubation, cells were harvested by centrifugation at 500 x g for 5 min, and cells were lysed in Pierce IP lysis buffer (Thermo Fisher) with HALT protease and phosphatase inhibitor cocktail (Thermo Fisher). Cells were allowed to lyse for 30 min on ice. Following incubation, lysates were clarified by centrifugation at 13, 000 x g for 10 min at 4 °C to pellet cell debris. Total protein was quantified using the detergent compatible Bradford reagent. Protein lysates were diluted across treatment groups to 1 mg/mL in 0.1X JESS sample buffer. Samples were separated using the Jess 12-230 kDa pre-filled plates (ProteinSimple, PS-PP03) and chemiluminescent module (ProteinSimple, PS-ST01EZ-8) as instructed by manufacturer, and blotted with the following antibodies: CARD9 (Santa Cruz, sc-374569) and GAPDH (Abcam, ab8245) or β-tubulin (CST, 2128) as loading controls.

### LDH Release Assay for Cytotoxicity Assessment

To assess cytotoxicity, cell supernatant was collected and mixed with LDH detection (Promega, J2380) enzyme mix and reductase substrate at 50 µL per reaction. Control wells received final concentration of 0.2% Triton X-100 to measure maximum LDH release. 300X sample dilution in LDH storage buffer was empirically determined using an assay linearity factor for LDH. Luminescence was recorded 30-60 min after the addition of LDH detection enzyme mix. Percent cytotoxicity was determined by the formula: % Cytotoxicity = [LDH activity of treated sample - LDH activity of control] / [(Total LDH activity) - (Control LDH activity)] × 100

### RT-qPCR Analysis

THP1 cells (100k/well) were plated in a 96-well plate and treated with 5 concentrations of CK1, CK2 inhibitors (10, 1, 0.1, 0.01, and 0.001 µM). Each sample was set in triplicates. The plates were incubated for 6 and 18 hr time points and centrifuged at 2000 rpm and the pellet was washed with PBS. Using NucleoSpin kit (Macherey-Nagel, 746709.4), pellets were lysed in 100 μL of RLT lysis buffer. Total RNA was extracted with on-column DNase treatment using manufacturer’s protocol and quantified using NanoDrop 8000 Spectrophotometer (Thermo Scientific). cDNAs were synthesized using 10 µL of total RNA from each sample with SuperScript™ IV VILO™ cDNA synthesis kit (Thermo Scientific, 11766050). PCR primers were used as shown in the below table. qPCR assay was set up using Ssofast Evagreen master mix (Bio-Rad, 1725203) in technical triplicates in a 384-well plate using manufacturer’s protocol and run on CFX384 Real time PCR machine.

Relative quantity was calculated using geometric mean of number of cycles of three house keeping markers – Ppia, Rpl13a and Actb using the following formulae: ΔΔCt = Ct (target gene) - Ct (Geomean) and RQ = 2-ΔΔCt. Fold change was calculated for each gene by normalizing RQ of sample with respective control ΔΔCq = RQ (sample)/RQ (DMSO Control).

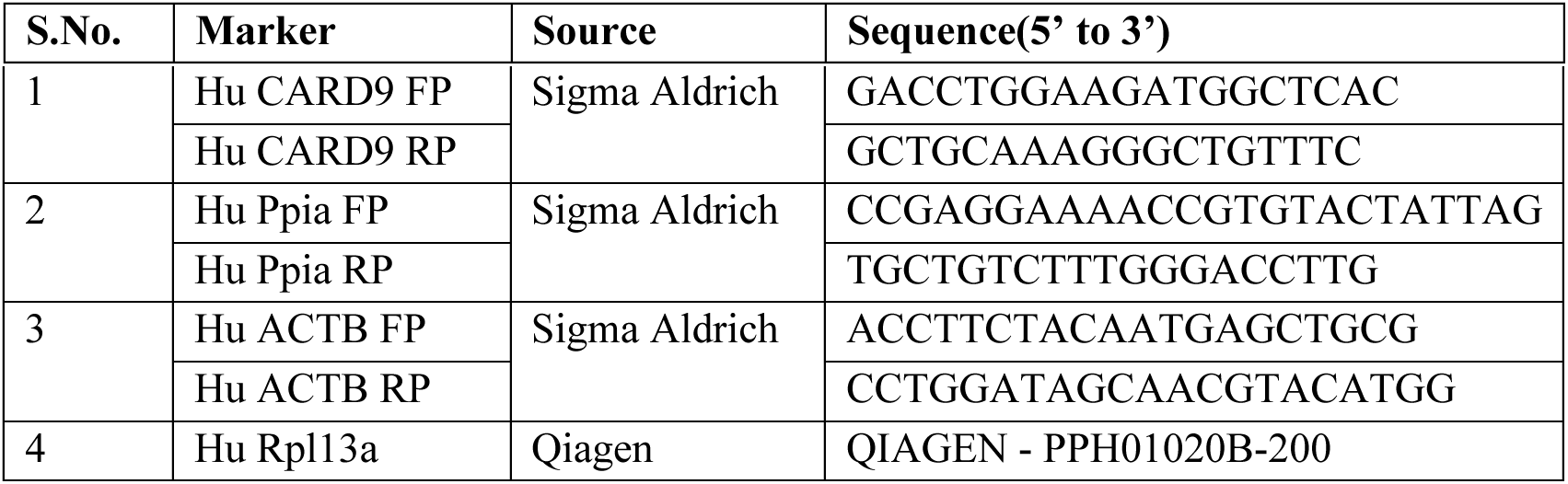

### Co-immunoprecipitation Experiment

Stable cell lines used in the co-immunoprecipitation experiments were generated by lentiviral transduction. For HEK293T FLAG-CARD9, lentivirus was made using Lentiviral Packaging Kit (Origene, TR30037) in HEK293T cells per manual’s instructions. Viral supernatants were harvested 48 hrs after transfections, and used to infect HEK293T cells in the presence of 20 µg/mL of polybrene. The cells were incubated with virus for 24 hrs before selection and validated via western blot.

To overexpress 3xFLAG-CK1α or CK2α, lentiviral expression plasmids were co-transfected with Lenti-X Packaging One Shot (Takara, 631276) into Lenti-X 293T cells, according to manufacturer’s protocol. Following 4 hr incubation fresh media was added to the cells. 48 hrs later viral supernatant was collected and filtered through a 0.45 µm cellulose acetate filter. HEK293T FLAG-CARD9 were later infected with CK1α or CK2α lentivirus supernatant using 5 μg/mL polybrene (Milipore Sigma). After 24 hrs, viral supernatant was removed and complete culture media was added to the cells containing 1 μg/mL puromycin. Cells were incubated for 48 hrs before selection. Cells were allowed to grow for an additional 4 days to select cells stably integrated with lentiviral expression plasmids. Expression of CARD9, CK1α and CK2α a was validated using JESS Western. Coding sequences of FLAG-CARD9, 3xFLAG-CK1α, 3xFLAG-CK2α a were synthesized by Genscript in a pLenti vector by Gibson assembly. All constructs were verified by Sanger sequencing.

Cell lysates were generated at 5 e^6^/mL in Pierce IP lysis buffer (Thermo Fisher) containing Halt protease and phosphatase inhibitor (Thermo Fisher). After cell pellets were harvested, washed and allowed to lyse on ice for 10 min in with periodic mixing, lysates were clarified at 13,000 x g for 10 min.

For the immunoprecipitation experiment following the CK2 inhibitor treatment, 2.5 e^6^ cells were plated on a 6-well plate and allowed to adhere in the assay media (DMEM with 10% HI-FBS). BMS-926 was added to the cells to achieve desired concentration (DMSO < 0.5%) and incubated at 37 °C. After 60 min, media containing the compound were aspirated and cells were washed with cold PBS and then lysed as described.

CK1α antibody (Abcam, ab206652), CK2α antibody (Santa Cruz, sc-373894) and its mouse IgG control (Santa Cruz, sc-2025) (4 μg for each sample) were complexed to Dynabeads Protein G magnetic beads (Thermo Fisher, 10007D) according to manufacturer’s instructions. Normalized lysates in 0.5 mL volume were incubated with antibody coated beads for 60 min at room temperature. After beads were washed with PBS, 30 μL 1X JESS sample master mix was added to the beads and boiled to elute the immunoprecipitated proteins, which were then visualized for CARD9 (Santa Cruz, sc-374569) and β-actin (Cell Signaling Technology, CST4970) on JESS as described above.

### Spectral Shift

MBP-CARD9 was obtained from Evotec. CARD9 was fluorescently labeled with the RED-NHS 2^nd^ Generation labelling kit (NanoTemper, MO-L011). The dye was reconstituted to 600 µM first then diluted further with labeling buffer supplied in the kit to a working solution of 100 uM. Ten microliter of this dye working solution was mixed with 20 μL CARD9 and 20 μL labeling buffer, and allowed to incubate for 2 hrs at room temperature in the dark.

After incubation, the reaction mixture was diluted with labeling buffer to make the final volume to 100 µL and added to the B-column to remove the excess dye. The B-column supplied in the kit was flushed 3 times with the CARD9 running buffer (25 mM HEPES, pH 7.3, 187 mM KCl, 3% glycerol, and 0.001% NP-40), then the diluted reaction mixture was added to the equilibrated B-column. After the sample fully entered the column bed, 450 µL of CARD9 running buffer was added and the flow through was collected. CARD9 running was buffer was added first at 500 µL and then 1000 µL to collect elution and post-wash elution respectively. Content of all tubes was analyzed at 330 and 350 nm using the NanoDSF (NanoTemper), with the highest signal in the elution fraction.

For the spectral shift assay between CK2 and CARD9, a fluorescent signal 8000 and 12000 RFU from labeled CARD9 was desired. Serial dilution of labeled CARD9 was analyzed on the Dianthus (NanoTemper) to determine dilution factor of the labeled CARD9 in the elution fraction.

For CK2 affinity measurement, a working solution of 200 µM CK2 (New England Bio, P6010L) and a working solution of 2x CARD9 (with 2x higher than the desired RFU) was prepared. ATP-gamma (Roche, 1162306001) concentration in each well was set at 100 µM. CK2 was titrated by adding 12 µL into the first well (final 100 µM) followed by 1:1 dilutions into CARD9 running buffer. 1 µL of ATP-gamma was added to each well and incubated for 15 min. Then 5 µL of labeled CARD9 was added to each well and allowed to incubate for 30 min before being read on the Dianthus. Data quality for spectral shift was assessed by 670/650 ratio greater than 0.007, the signal-to-noise being greater than 5, and the saturation of the curve being greater than 90%.

### Biochemical Phosphorylation

12 μM MBP-CARD9 and 8 μM CK2 were mixed in 20 µL of reaction buffer (25 mM HEPES, pH 7.3, 187 mM KCl, 3% glycerol, and 0.001% NP-40) in the presence of 100 μM ATP (Roche) and the mixture was incubated at room temperature for 2 hrs. The reaction mixture was mixed with PreOmics BCT-LYSE and tryptic lysates were further prepared for proteomic data acquisition using the PreOmics iST-BCT 96 sample kit. Protein digests were subjected to data collection using an LC-MSMS system (Evosep, #EV-1000) connected online to a mass spectrometer (Bruker, timsTOF pro2) in data dependent acquisition mode. Data was searched against the fasta file containing experiment specific MBP-CARD9 and CK2 using the software MaxQuant v2.1.3.0 with the following parameters: Number of modifications was maxed at 5 per peptide and included Cys Carbamidomethylation as fixed modification, and phosphorylation (STY), Ox(Met) and acetylation on protein N-term as variable modification. Missed cleavages were set to max 2. Minimum peptide length is 5 and with a 1.0% FDR for PSMs and proteins. MS/MS tolerance was set to 25 ppm, and match between runs was enabled.

### NanoBRET Assays

Coding sequences of Halo and NanoLuc tagged CARD9 and CK2 were synthesized by Genewiz and cloned into the pCDH-UbC-MCS-EF1α-Puro vector (System Biosciences) by Gibson assembly. All constructs were verified by Sanger sequencing. To evaluate protein-protein interaction of CARD9 and CK2, HEK-293T cells (1.5 e^6^) were transiently transfected with 1 µg tDNA at a 1:100 ratio of NL:HT vectors. CK2-NanoLuc was co-expressed with HaloTag-CARD9, HaloTag-CARD9_T231D, or HaloTag-CARD9_del11 in 6-well plates using Lipofectamine 2000 (Invitrogen). After 48 hrs, the transfected cells were harvested and resuspended in fresh assay medium (Opti-MEM reduced serum, no phenol red with 4% HI-FBS, Gibco) and plated at 32k cells/ 50 µL in a 96-well plate. Using a Tecan, cells were dosed with compounds in the absence or presence of 0.1 mM HaloTag NanoBRET 618 Ligand (Promega, N1661) and incubated for 4 hrs at 37° C. NanoBRET Nano-Glo Substrate was prepared in Opti-MEM Reduced Serum Medium, no phenol red at a 100-fold dilution. 12 µL of solution was added to each well and spun at 600 x g for 15 sec. Donor emission (460 nm) and acceptor emission (618 nm) was measured immediately on GloMax (Promega). BRET ratio is calculated by dividing acceptor emission by donor emission. Percent inhibition was determined by the formula: % inhibition = [BRET ratio of treated sample with NanoBRET ligand – BRET ratio in absence of the ligand] / [(BRET ratio of DMSO sample with NanoBRET ligand – BRET ratio in absence of NanoBRET ligand)] × 100

### Phosphoproteomics

THP-1 cells were seeded in 96-well plates and treated with the compound for the indicated time. Tryptic lysates were prepared for proteomic data acquisition using the PreOmics iST-BCT 96 sample kit. Phosphopeptide samples were then enriched using Agilent Bravo with Fe(III)-NTA cartridges tips (Agilent, G5496-60085) followed by data collection using an LC-MSMS system (Evosep, #EV-1000) connected online to a mass spectrometer (Bruker, timsTOF SCP). Peptide identification and relative quantification were performed using DIA-NN v1.8.1^2^ with an experiment specific library generated from THP-1 cells. Precursor data obtained from DIA-NN was re-summarized with using the tidyverse package. Briefly, each peptide was grouped by its root sequence, charge state, number of phosphorylations, and protein group to create unique precursor group identifiers, and the quantitative data were summarized by calculating the mean across observations within each group. The resummarized data was then subjected for imputation with using Bayesian Principal Component Analysis (BPCA) via the pcaMethods package^5^ and a minimal detection imputation (MinDet) via the imputeLCMD package^3^ for low-abundance features, setting a quantile threshold at 0.01. Differential expression analysis was performed on the imputed data with using the limma package.^4^

### Computational Modeling

Since there is no publicly available full-length CARD9 structure, the predicted monomeric structure of CARD from AlphaFold^6–8^ was used for CARD9 along with the kinase domain of CK2 from 5ZN1^9^ structure from RCSB PDB. Both structures were prepared using Protein Preparation Wizard in the Schrödinger software suite (Release 2023-1) prior to protein-protein docking.^10^ Protein-protein docking was carried out using Piper (Schrödinger)^11–13^ with distance constraints between S424/S425 from CARD9 and ATP binding domain of CK2. The protein-protein interfaces (PPIs) of the resulting 30 poses of CARD9:CK2 complex from protein-protein docking were refined using Prime (Schrödinger)^14–15^ to remove clashes and to calculate energies. Interaction fingerprints of PPIs were calculated using Canvas^16–18^, which were then used to cluster the poses into five clusters. The centroids from three clusters that have more than five members were selected to evaluate the stability of the PPIs.

Two molecular dynamics (MD)-based methods were used for assessing the stability using Desmond MD engine in the Schrödinger Suite. First, three replicates of plain MD simulations were carried out for each cluster centroid poses of CARD9:CK2 complex. Each system was first solvated in 12 Ang orthorhombic box of SPC water molecules with neutralizing ions. The system was heated to 10 K with restraints on solute heavy atoms for 100 ps in Brownian Dynamics in NVT ensemble, followed by MD simulation in NVT ensemble for 12 ps. Then MD simulations were run in NPT ensembles for 12 ps each with and without restraints on solute heavy atoms while heating up to 300 K. Finally, the production simulations were run at 300 K for 100 ns. Different random seed for initial velocity was used for each of three replicates. After the simulations were finished, root mean square deviation (RMSD) and root mean square fluctuation (RMSF) of CK2, or “ligand” heavy atoms were calculated with respect to the given starting pose to assess the stability of the PPI.

Well-tempered metadynamics was used as another measure of stability estimation for PPIs. The same equilibration procedure as above for the plain MD simulations was carried out. For production stage, well-tempered metadynamics was performed by applying biasing potentials to the selected collective variables (CVs) to flatten the free energy surface of interest. The backbone atoms of the residues within 4 Å and 6 Å of S424 and S425 of CARD9 and CK2, respectively, were chosen as CVs. Three replicates of simulations were carried out for each system with different random seeds for initial velocities. After the simulations, average RMSD from the starting pose was calculated across three replicates. The PPI with lowest average RMSD was selected to be the most stable system.

### Dual-Luciferase Reporter Assay for Phosphomimetics screen

In a 96-well format, 25k HEK293T cells were co-transfected using Lipotectime 3000 (Invitrogen) according to the manufacturer’s instructions with pNF-κB luciferase reporter, pRL-TK *Renilla* Reporter, CARD9 and BCL10 plasmids normalized to 175 ng of plasmid per well. Following a 20 hr incubation at 37 °C, firefly luciferase activity was measured using the Dual-luciferase reporter assay system (Promega). This was followed by measuring *Renilla* luciferase activity to normalize transfection efficiencies. Coding sequences of CARD9 and BCL10 were synthesized by Genscript in a pLenti vector by Gibson assembly. All constructs were verified by Sanger sequencing.

### Western Blot

HEK293T cells were harvested 20 hr following transfection and lysed in Pierce IP Lysis Buffer (Thermo Fisher) with cOmplete™ Protease Inhibitor Cocktail (Roche). Protein concentrations were determined by Bradford Assay. 10 μg of protein was prepped with 6x Laemmli SDS sample buffer (Thermo Fisher) and separated in 4-12% Bis-Tris gels (Invitrogen). The gels were transferred to Nitrocellulose membranes using iBlot™ Transfer Stack (Thermo Fisher). Membranes were incubated at 4 °C overnight with the following antibodies: 1:500 anti-FLAG (Cell Signaling Technology, CST14793), 1:2000 anti-GAPDH (Abcam, ab8245). Blots were visualized using LI-COR Odyssey Image System.

### Native Gel Electrophoresis

Cell lysates were made using the NativePAGE™ Sample Prep Kit (Invitrogen) according to instructor’s manual. Protein concentrations were determined by Bradford Assay (ThermoFisher) and 10 ug of protein was separated in NativePAGE 4-12% Bis-Tis gels (Invitrogen). The gels were transferred to PVDF membranes using iBlot™ Transfer Stack (Thermo Fisher). The membranes were incubated at 4 °C overnight with 1:1000 anti-FLAG (Cell Signaling Technologies, CST14793). Blots were visualized using LI-COR Odyssey Image System.

### CARD9 Quantification in Tissue Homogenate via AlphaLISA

In preparation for the peritonitis model with the CK2 inhibitor, mouse AlphaLISA kit was validated first with CARD9^WT^ and CARD9^KO^ mice to quantify CARD9 levels in tissue homogenates. The model and tissue harvesting protocol is described above. Subsequent mouse CARD9 AlphaLISA was performed according to manufacturer’s instructions.

Fresh whole blood samples were centrifuged at 4000 rpm for 10 min to extract plasma. The samples were washed with 400 µL of PBS, and a 200 µL aliquot of the blood/PBS suspension was transferred into 96-well round-bottom plates and centrifuged at 2000 rpm for 5 min. The PBS was aspirated, and the remaining pellet was lysed in 200 µL of 10x eBioscience Buffer, diluted to 1x with deionized water, for 3 min. The plates were centrifuged at 600 x g for 3 min, and the supernatant was removed. A second lysis was performed with 200 µL of 1x eBioscience Buffer for 2.5 min, followed by centrifugation. The supernatant was aspirated, and all wells were washed with 200 µL of dPBS. The washed pellets were lysed in 50 μL 1x CARD9 AlphaLISA Lysis Buffer for 45 min in cold room on shaker. Samples were clarified by centrifugation following lysis and protein concentration was normalized to 1 mg/mL.

Peritoneal lavage samples were received in PBS suspension in 500 µL deep-well 96-well plates. The plates were centrifuged at 500 x g for 5 min, and the supernatant was aspirated. The remaining pellets were lysed in 25 µL of 1x CARD9 AlphaLISA Lysis Buffer for 45 min on a shaker in a cold room. Following lysis, the samples were clarified by centrifugation, and the protein concentration was normalized to 0.25 mg/mL.

Frozen colon samples were received in Precellys tubes. Each tube was supplemented with 1 mL of CARD9 AlphaLISA Buffer and subjected to homogenization three times. The homogenates were then incubated on ice for 20 min to allow for lysis. Following lysis, the samples were clarified by centrifugation. The protein concentration of the clarified lysates was normalized to 1.5 mg/mL.

## Chemical Synthesis and Spectra

### Chemical compound used in this study

BMS-926,^18^ BMS-713,^19^ BMS-747^19^ were previously prepared, and preparation of BMS-321 is described below. CX-4945 was purchased from MedChemExpress.

### Synthesis Procedure and Spectra

**Preparation of N-(6-(3-(4-fluorophenyl)-1-methyl-1H-pyrazol-4-yl)-[1,2,4]triazolo[1,5-a]pyridin-2-yl)benzamide** (**BMS-321**).

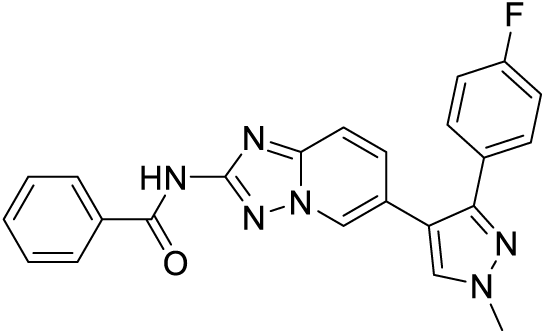

**<u>Step 1:</u> Preparation of 6-(3-(4-fluorophenyl)-1-methyl-1H-pyrazol-4-yl)-[1,2,4]triazolo[1,5-a]pyridin-2-amine.**

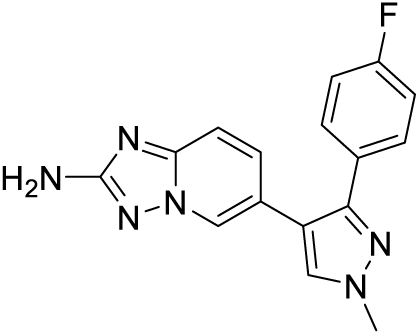

A mixture of 6-bromo-[1,2,4]triazolo[1,5-a]pyridin-2-amine (115 mg, 0.540 mmol), 3-(4-fluorophenyl)-1-methyl-4-(4,4,5,5-tetramethyl-1,3,2-dioxaborolan-2-yl)-1H-pyrazole (428 mg, 0.567 mmol) (previously prepared^20^) and 1,1’-Bis(di-tert-butylphosphino)ferrocene palladium dichloride (35.2 mg, 0.054 mmol) was degassed. Dioxane (2.7 mL) was added followed by potassium phosphate (540 µL, 1.62 mmol), and the reaction mixture was degassed, purged with nitrogen, and heated at 90 °C overnight. The reaction mixture was cooled and then filtered through a pad of celite, and the filtrates were partitioned between water and ethyl acetate. The organic phase was washed with brine, dried over anhydrous magnesium sulfate, filtered, and concentrated under reduced pressure. The viscous product mixture was dissolved in dimethylformamide and purified by reverse phase, preparative HPLC to give 6-(3-(4-fluorophenyl)-1-methyl-1H-pyrazol-4-yl)-[1,2,4]triazolo[1,5-a]pyridin-2-amineas a white solid (43 mg).

^1^H NMR (499 MHz, chloroform-d) δ 8.23 - 8.19 (m, 1H), 7.50 (s, 1H), 7.45 (dd, J=8.9, 5.4 Hz, 2H), 7.34 (dd, J=9.0, 0.8 Hz, 1H), 7.25 (dd, J=9.0, 1.8 Hz, 1H), 7.03 (t, J=8.7 Hz, 2H), 4.46 (s, 2H), and 4.00 (s, 3H).

**<u>Step 2:</u> Preparation of N-(6-(3-(4-fluorophenyl)-1-methyl-1H-pyrazol-4-yl)-[1,2,4]triazolo[1,5-a]pyridin-2-yl)benzamide (BMS-321).**

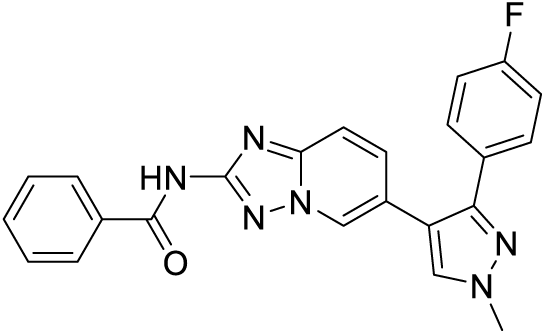

To a solution of 6-(3-(4-fluorophenyl)-1-methyl-1H-pyrazol-4-yl)-[1,2,4]triazolo[1,5-a]pyridin-2-amine (42 mg, 0.136 mmol) and Hunig’s base (71.4 µL, 0.409 mmol) in dichloromethane (2 mL) at room temperature was added a solution of benzoyl chloride (17.4 µL, 0.150 mmol) in dichloromethane (252 µL) dropwise. The resulting reaction mixture was stirred at room temperature for 2 h. Additional Hunig’s base (71.4 µL, 0.409 mmol) and additional benzoyl chloride (17.4 µL, 0.150 mmol) were added. After additional 2h, the reaction mixture was partitioned between 1.5 M K_3_PO_4_ and ethyl acetate. The organic phase was washed with brine, dried over magnesium sulfate, filtered under reduced pressure, and concentrated. The resulting residue was dissolved in dimethylformamide and purified by reverse phase, preparative HPLC to give N-(6-(3-(4-fluorophenyl)-1-methyl-1H-pyrazol-4-yl)-[1,2,4]triazolo[1,5-a]pyridin-2-yl)benzamide (**BMS-321**) as a tan solid (17 mg).

HPLC purity: 99% (t_r_ = 0.905 min. - Column: Waters Acquity BEH C18, 2.1 x 50 mm, 1.7-μm particles; Mobile Phase A: 5:95 acetonitrile:water with 0.05% trifluoroacetic acid; Mobile Phase B: 95:5 acetonitrile:water with 0.05% trifluoroacetic acid; Temperature: 50 °C; Gradient: 0-100% B over 1.0 min, then a 0.5 min hold at 100% B; Flow: 1.0 mL/min; Detection: UV at 220/254 nm);

LCMS (ESI) m/z Calcd for C_23_H_17_FN_6_O [M + H]+ 413.2. Found: 413.1.

^1^H NMR (499 MHz, Methanol-d4) δ 8.56 (d, J=0.7 Hz, 1H), 8.03 - 7.99 (m, 2H), 7.95 (s, 1H), 7.66 - 7.58 (m, 2H), 7.57 - 7.46 (m, 5H), 7.12 (t, J=8.8 Hz, 2H), and 4.00 (s, 3H).

^13^C NMR (126 MHz, DMSO-d6) δ 165.5, 163.2, 161.3, 159.2, 148.7, 147.2, 134.2,132.6, 132.2, 130.4, 130.3, 129.9, 128.9, 128.9, 128.5, 128.5, 126.8, 120.2, 116.1, 115.9, 115.2, 115.1, 39.3.

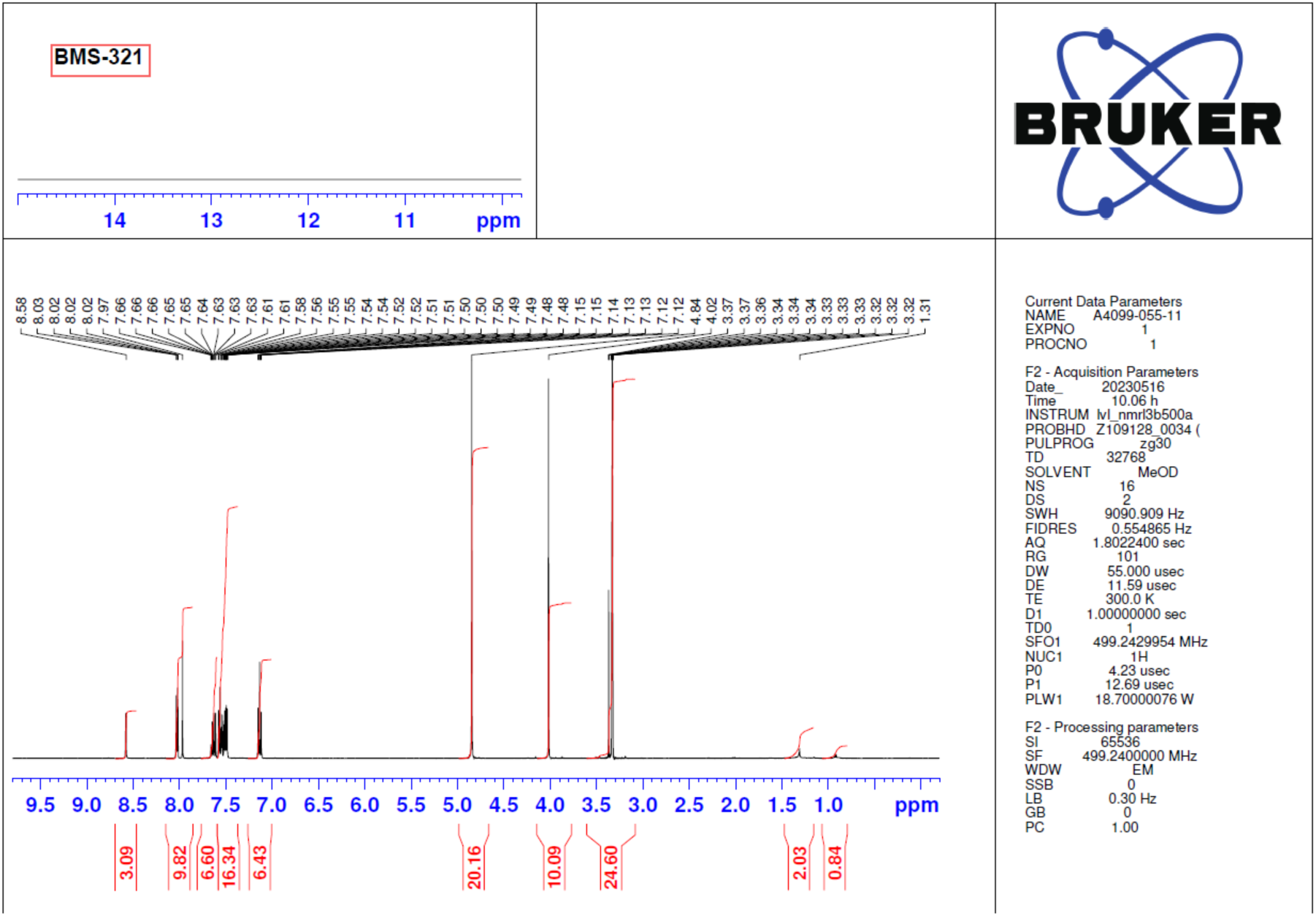

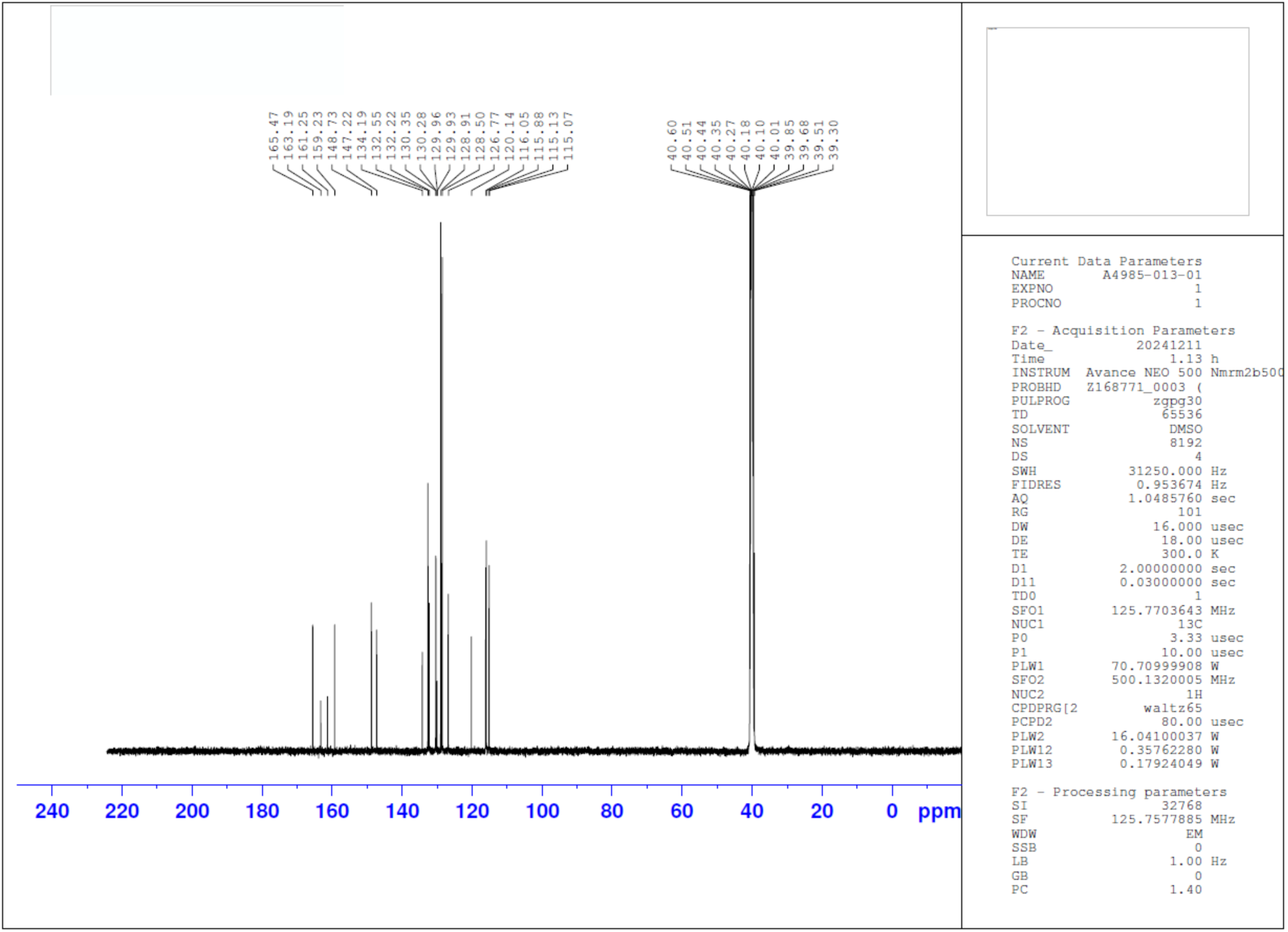

